# Multiscale profiling of enzyme activity in cancer

**DOI:** 10.1101/2021.11.11.468288

**Authors:** Ava P. Soleimany, Jesse D. Kirkpatrick, Cathy S. Wang, Alex M. Jaeger, Susan Su, Santiago Naranjo, Qian Zhong, Christina M. Cabana, Tyler Jacks, Sangeeta N. Bhatia

## Abstract

Diverse processes in cancer are mediated by enzymes, which most proximally exert their function through their activity. Methods to quantify enzyme activity, rather than just expression, are therefore critical to our ability to understand the pathological roles of enzymes in cancer and to harness this class of biomolecules as diagnostic and therapeutic targets. Here we present an integrated set of methods for measuring specific enzyme activities across the organism, tissue, and cellular levels, which we unify into a methodological hierarchy to facilitate biological discovery. We focus on proteases for method development and validate our approach through the study of tumor progression and treatment response in an autochthonous model of *Alk*-mutant lung cancer. To quantitatively measure activity dynamics over time, we engineered multiplexed, peptide-based nanosensors to query protease activity *in vivo*. Machine learning analysis of sensor measurements revealed dramatic protease dysregulation in lung cancer, including significantly enhanced proteolytic cleavage of one peptide, S1 (*P_adj_* < 0.0001), which returned to healthy levels within three days after initiation of targeted therapy. Next, to link these organism-level observations to the *in situ* context, we established a multiplexed assay for on-tissue localization of enzyme activity and pinpointed S1 cleavage to endothelial cells and pericytes of the tumor vasculature. Lastly, to directly link enzyme activity measurements to cellular phenotype, we designed a high-throughput method to isolate and characterize proteolytically active cells, uncovering profound upregulation of pro-angiogenic transcriptional programs in S1-positive cells. Together, these methods allowed us to discover that protease production by angiogenic vasculature responds rapidly to targeted therapy against oncogene-addicted tumor cells, identifying a highly dynamic interplay between tumor cells and their microenvironment. This work provides a generalizable framework to functionally characterize enzyme activity in cancer.

## Introduction

Nearly all processes in malignant progression rely on dynamic changes in the activity, not just abundance, of biomolecules. Methods to quantitatively track protein activity within the cellular, spatial, and organismic contexts are therefore critical to achieve a systems-level understanding of cancer biology and to design next-generation precision cancer medicines. However, while the omics revolution has enabled high-throughput assays of the genome, epigenome, transcriptome, and proteome [1], it has largely stopped short of queries at the protein activity level–a distinct axis of regulation that is often most proximal to actuated biological function. Although single-cell transcriptomics has enabled characterization of intratumoral heterogeneity [2, 3, 4], and techniques to localize proteins [5, 6] and nucleic acid sequences [7, 8, 9] *in situ* are starting to enable study of tumors in a spatial context [10], analogous techniques for single-cell and spatial profiling of protein activity have been largely undeveloped. To achieve a more comprehensive understanding of cancer pathophysiology across the organism, tissue, and cellular scales, methods to directly quantify tumor-associated protein activity in these contexts are required.

Recent years have seen a push to develop biosensors that measure biomolecular activity *in vivo* to generate synthetic signals that can be read out noninvasively [11, 12, 13, 14, 15, 16]. For example, *in vivo* sensors of enzyme activity have enabled noninvasive detection of cancer [11, 12, 17, 18, 19, 20], while active glucose uptake has been used for functional imaging of cancer metabolism [21]. However, such *in vivo* readouts have largely treated the body as a black box, sacrificing crucial information on spatial localization within the tumor microenvironment (TME), precluding dissection of phenotypic heterogeneity at the single-cell level, and thus reducing meaningful biological interpretation. Therefore, there remains a need for methods capable of generating and unifying molecular activity measurements across biological scales.

In this work, we present an integrated set of methods to profile enzyme activity in cancer across the organism, tissue, and cellular scales (Fig. 1). As an initial target for method development, we focus on proteases, enzymes that are dysregulated in cancer and directly contribute to all cancer hallmarks [22]. In the *in vivo* setting, we leverage multiplexed protease-responsive nanosensors together with machine learning to noninvasively and longitudinally monitor activity dynamics within living organisms. To explore tissue-level organization within the TME, we establish a multiplexed assay for on-tissue spatial localization of protease activity against target peptide substrates. Finally, to link protease activity to other measurement modalities at the cellular scale, we design a single-cell method, termed activity-based cell sorting, that uses peptide probes and flow cytometry to sort individual cells based on their associated enzymatic activity.

**Figure 1:**
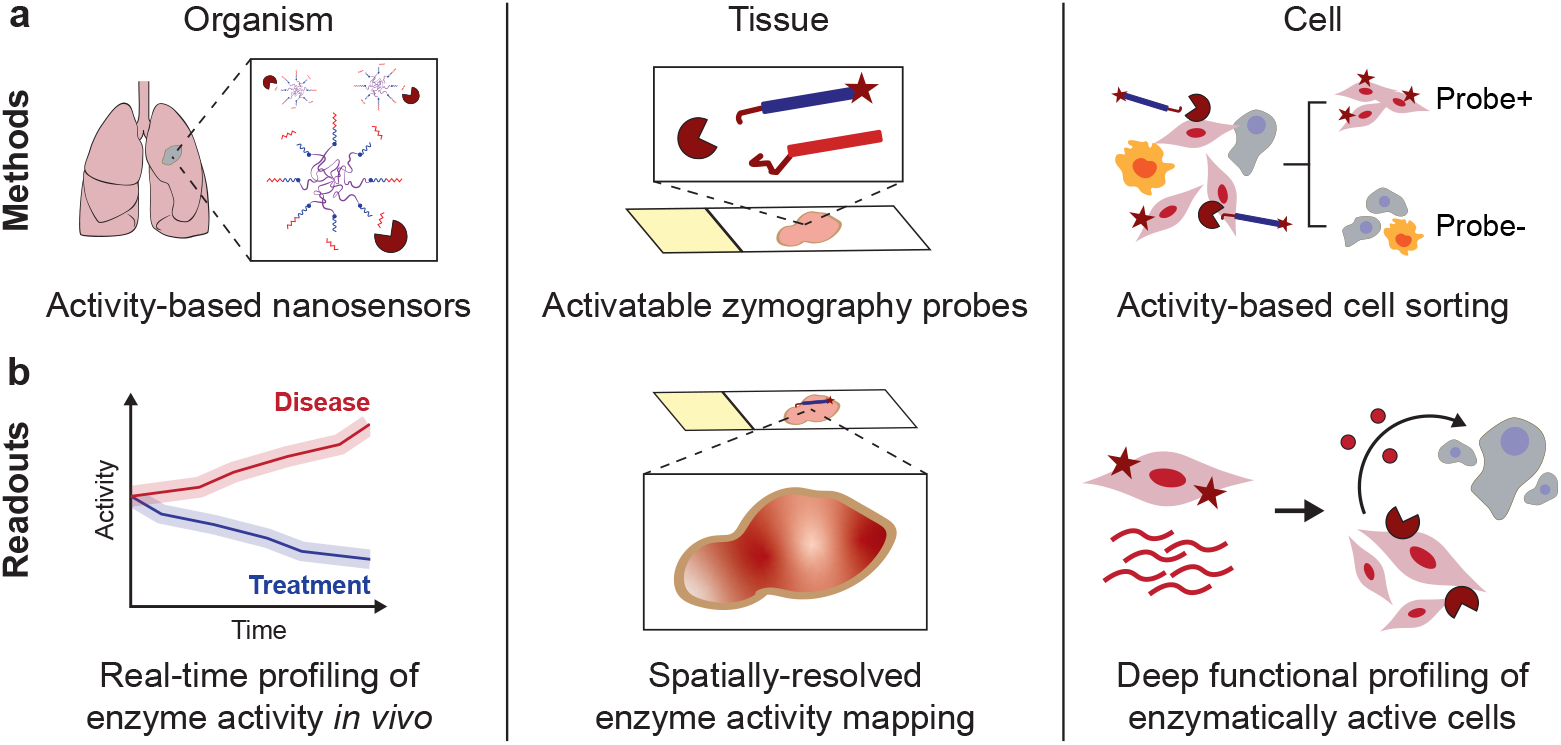
Multiscale profiling of enzyme activity in cancer. **(a)** Methods for profiling enzyme activity across the organism, tissue, and cellular scales. Noninvasive activity-based nanosensors can be translated into activatable probes for *in situ* zymography and activity-based cell sorting. **(b)** Method readouts enable noninvasive, real-time monitoring of *in vivo* enzyme activity over tumor progression and treatment response; high-resolution, spatially-resolved enzyme activity mapping of the TME; and single-cell isolation and multimodal characterization of enzymatically active cells.

We unified these methods into a single hierarchical framework to power biological discovery (Fig. 1), and applied it to study tumor progression and early drug response in an autochthonous mouse model of *Alk*-mutant non-small-cell lung cancer (NSCLC) [23]. Using this framework, we uncovered profound shifts in protease activity that occur after targeted therapy with the ALK inhibitor alectinib. Spatial and single-cell profiling linked a treatment-responsive activity signature to pericytes and endothelial cells of the angiogenic tumor vasculature. Our results reveal a dynamic cross-talk between cancer cells and cells of the TME; demonstrate that protease activity can serve as a powerful proxy for specific cancer hallmarks; and validate the utility of multiscale enzyme activity profiling for functional characterization of cancer biology.

## Results

### Profiling enzyme activity *in vivo* to noninvasively monitor tumor progression

We first sought to establish the ability of our activity-based profiling framework to noninvasively quantify changes in biological activity as a function of tumor progression and treatment response. We utilized an autochthonous mouse model of ALK^+^ NSCLC as a model system, in which intrapulmonary administration of an adenovirus encoding two guide RNAs and Cas9 resulted in oncogenic rearrangement of the *Eml4* and *Alk* genes (the “Eml4-Alk” model; Fig. 2a), leading to the formation of lung tumors that histologically resembled human lung adenocarcinoma [23]. We queried a bulk RNA sequencing (RNA-seq) dataset of Eml4-Alk lungs [24] and identified several proteases overexpressed in Eml4-Alk mice (Fig. S1). To noninvasively monitor protease activity in the Eml4-Alk model, we engineered a multiplexed panel of activity-based nanosensors that can be selectively activated by dysregulated proteases within the TME to release barcoded reporters into the urine [19]. Critically, these nanosensors (Table S1) carry peptide substrates that can be recognized *in vitro* by a range of metallo-, serine, and aspartic proteases [19], and require substrate cleavage for activation and barcode release. At 3.5 weeks after tumor induction, we intratracheally administered the nanosensor panel into Eml4-Alk and healthy control mice, and observed that several nanosensors were differentially cleaved by proteases in the pulmonary microenvironment (Fig. S2a), enabling robust separation of the two groups (Fig. S2b).

**Figure 2:**
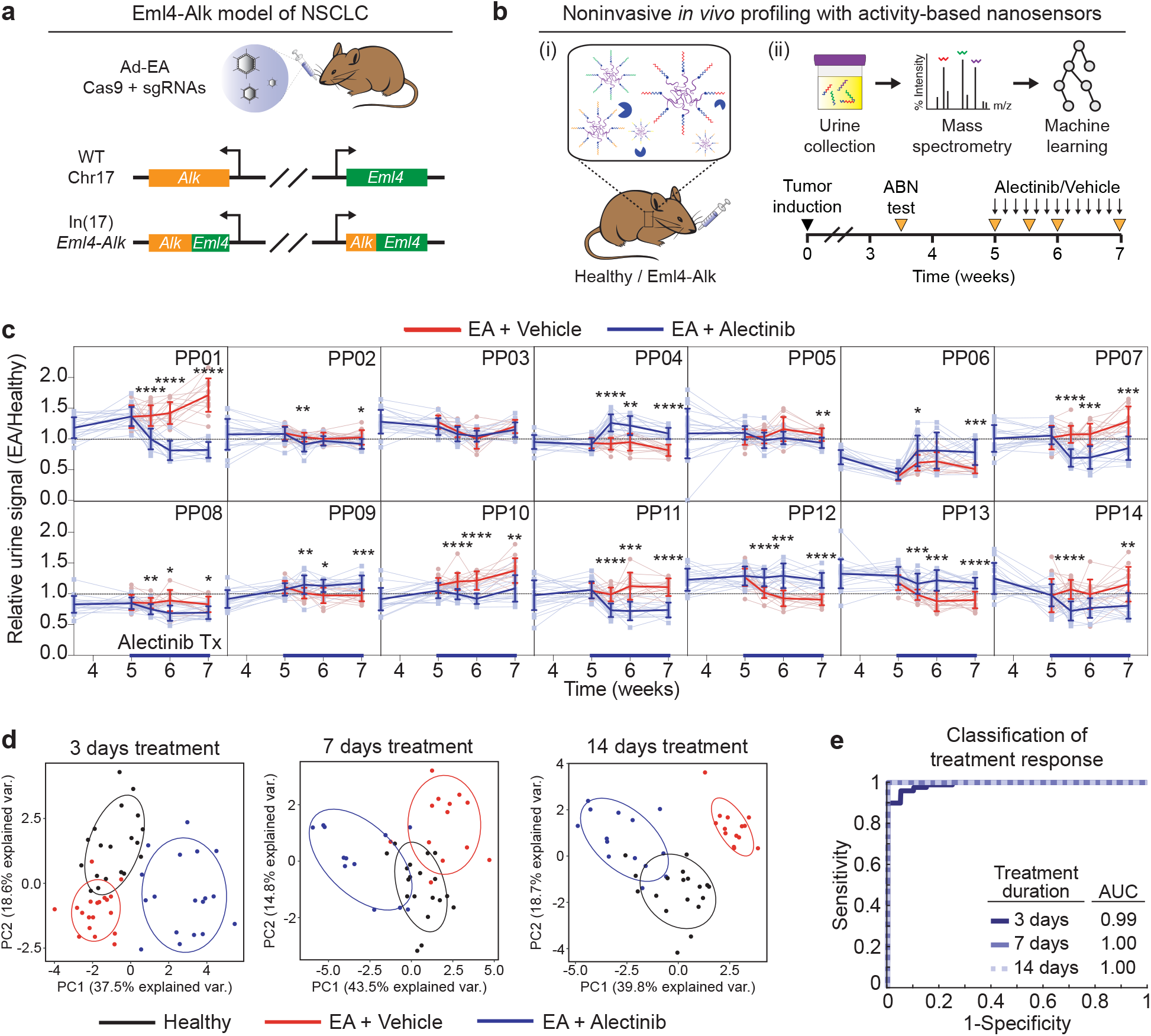
Activity-based nanosensors measure *in vivo* enzyme activity dynamics over tumor progression and treatment response. **(a)** Disease was induced in the Eml4-Alk model via CRISPR-Cas9 mediated rearrangement of the *Eml4* and *Alk* loci. **(b)** Schematic of approach. (i) Activity-based nanosensors were administered to Eml4-Alk and healthy mice by intratracheal instillation. (ii) Tumor-bearing Eml4-Alk mice were treated with either alectinib or vehicle and subject to *in vivo* protease activity profiling (ABN test) over the course of disease progression. **(c)** Healthy control-normalized urinary reporter signal for each of the 14 activity-based nanosensors. Transparent lines show trajectories of each mouse over time; opaque lines are averages over all mice per group. Red lines represent Eml4-Alk mice treated with vehicle (EA + Vehicle; n = 20, 19, 13, and 14 for 5, 5.5, 6, and 7 weeks, respectively), and blue lines represent Eml4-Alk mice treated with alectinib (EA + Alectinib; n = 20, 19, 12, and 14 for 5, 5.5, 6, and 7 weeks, respectively), with n = 20 at 3.5 weeks (mean ± s.d; multiple t-tests with Holm-Sidak correction; \**P* < 0.05, \*\**P* < 0.01, \*\*\**P* < 0.001, \*\*\*\**P* < 0.0001. **(d)** Principal component analysis of mean scaled urinary reporter concentrations of healthy (Healthy), vehicle-treated Eml4-Alk (EA + Vehicle), and alectinib-treated Eml4-Alk (EA + Alectinib) mice at 3, 7, and 14 days post treatment induction. **(e)** Receiver operating characteristic (ROC) curve performance of a random forest classifier, trained on urinary reporters from Eml4-Alk mice, in discriminating an independent test cohort at the designated post-treatment time points (n = 10 independent trials).

We then assessed whether activity-based nanosensors could rapidly and quantitatively monitor the dynamics of tumor progression and regression. We treated Eml4-Alk mice with the first-line clinical ALK inhibitor alectinib [25] and monitored changes in pulmonary protease activity over a two-week treatment course that resulted in rapid and robust tumor regression (Fig. 2b, Fig. S3a-b). Strikingly, we observed that alectinib treatment dramatically altered pulmonary protease activity within just 3 days of treatment initiation, with 12 of 14 reporters exhibiting significantly differential enrichment in the urine of vehicle-versus alectinib-treated mice (Fig. S4). Signal trajectories for each of the individual sensors revealed several distinct patterns in their dynamics (Fig. 2c). Notably, cleavage of a subset of sensors (e.g., PP01, PP07, PP10) increased over time in vehicle-treated mice as tumors progressed but rapidly regressed following alectinib treatment, while the cleavage of other sensors (e.g., PP04) transiently increased upon initiation of alectinib treatment and then returned towards baseline levels. Collectively, these measurements revealed increased divergence in proteolytic activity signatures from vehicle-treated Eml4-Alk mice versus healthy controls over the course of tumor progression, while signatures from alectinib-treated Eml4-Alk mice exhibited greater similarity to those of healthy controls as a function of treatment (Fig. 2d). Finally, a random forest classifier trained on urinary reporter signatures from a subset of Eml4-Alk mice achieved highly accurate classification of therapeutic response to ALK inhibition (Fig. 2e). These results validate a method that quantitatively measures enzyme activity in living organisms and thus enables noninvasive, real-time monitoring of cancer progression.

### Multiplexed spatial localization of enzyme activity

We next sought to establish a method to mechanistically explore the biological drivers of activity signatures identified by our *in vivo* platform. To this end, we reasoned that tissue-level spatial localization of enzyme activity against target substrates could facilitate biological interpretation. Because our *in vivo* nanosensors use peptide cleavage as their measurement mechanism, we directly translated their substrates into *in situ* activatable zymography probes (AZPs) that also rely on substrate-specific proteolytic cleavage for activation [26]. Within an AZP, a protease-cleavable substrate links a fluorophore-tagged, positively-charged domain (polyR) with a negatively-charged domain; this structure remains complexed in the absence of proteolytic activation. When AZPs are applied to fresh-frozen tissue sections in a manner analogous to immunofluorescence staining, substrate cleavage by tissue-resident enzymes liberates the tagged polyR to electrostatically interact with and bind the tissue, enabling localization of protease activity by microscopy.

We thus leveraged AZPs for on-tissue spatial localization of protease activity against target peptide substrates of interest, i.e., those nominated from our *in vivo* activity screen. We selected three nanosensors whose signal tracked with tumor progression and responded to alectinib treatment (PP01, PP07, PP10; Fig. 2c), and incorporated them into individual AZPs with orthogonal fluorophores (Z1, Z7, Z10, respectively; Fig. 3a). We reasoned that a multiplexed solution of these AZPs would allow for simultaneous profiling and localization of proteolytic cleavage of each of these three substrates. We applied the three-plex AZP solution to fresh-frozen lung tissue sections from Eml4-Alk mice at 7 weeks post tumor induction (Fig. 3b) and detected fluorescence signal from each probe within Eml4-Alk tumors (Fig. 3c-d). Strikingly, this multiplexed *in situ* labeling revealed a differing pattern of spatial localization for Z1 relative to Z7 and Z10 (Fig. 3d). Qualitatively, while Z7 and Z10 exhibited broad, diffuse staining throughout Eml4-Alk tumor tissue, Z1 exhibited a prominent spindle-like pattern, suggesting that cleavage of each of those probes could correspond to distinct proteases and biological processes (Fig. 3d). Tissue labeling by each of the three probes was significantly abrogated by addition of a cocktail of protease inhibitors, indicating successful multiplexing of multiple AZPs (*P* < 0.0001; Fig. 3e-g).

**Figure 3:**
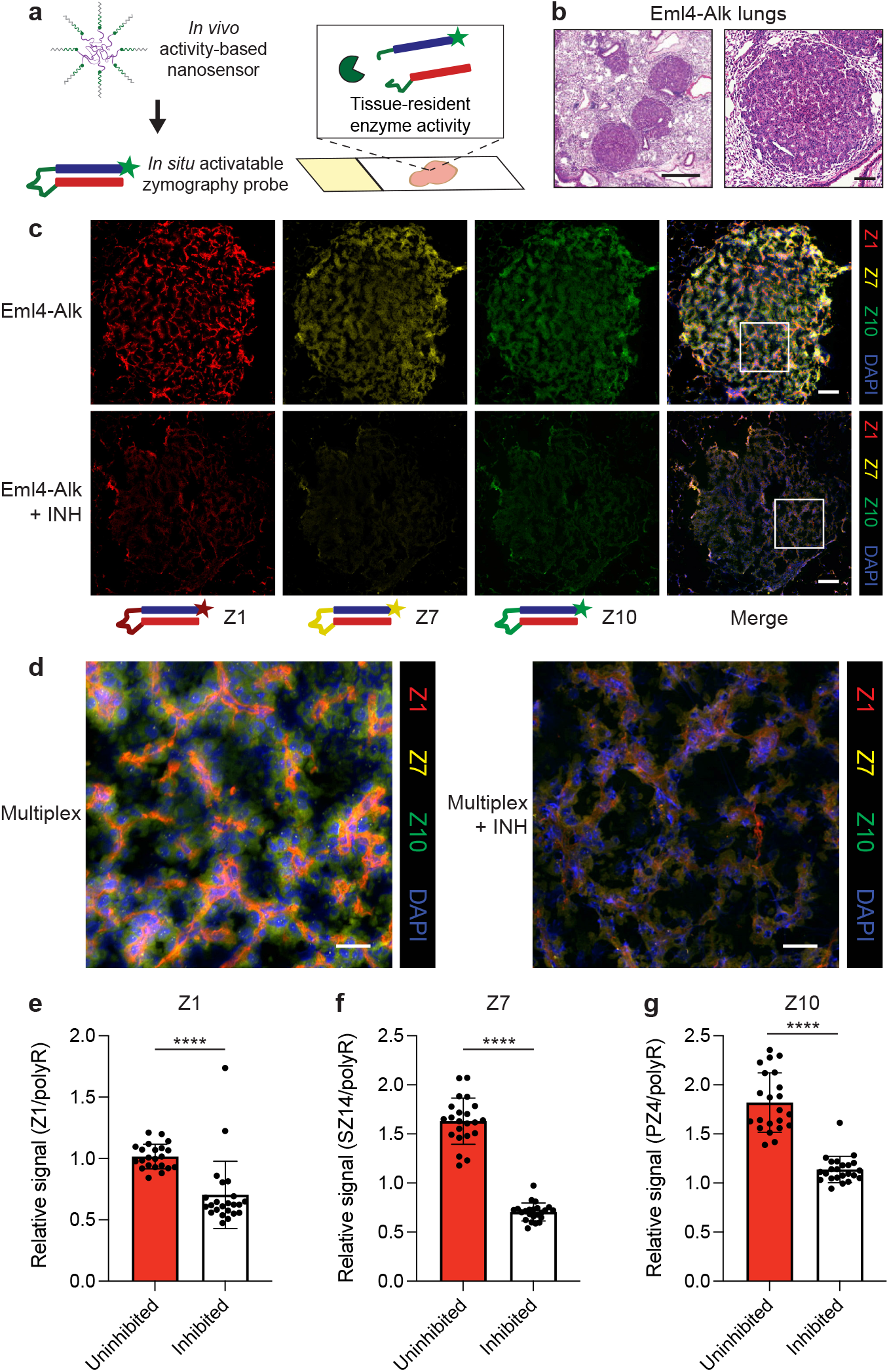
Multiplexed spatial localization of protease activity with AZPs. **(a)** Substrates nominated from *in vivo* profiling were translated into *in situ* AZPs to measure and spatially localize tissue-resident enzyme activity in frozen tissue sections. **(b)** AZPs were applied to fresh-frozen lung tissue sections from Eml4-Alk tumor-bearing mice. Haematoxylin and eosin (H&E) stains of representative Eml4-Alk tumors. Scale bars = 500 *μ*m (left), 100 *μ*m (right). **(c)** Application of a multiplexed AZP cocktail of Z1 (red), Z7 (yellow), and Z10 (green), with or without broad-spectrum protease inhibitors, to Eml4-Alk lung tissue sections (Eml4-Alk, Eml4-Alk + INH, respectively). Scale bars = 100 *μ*m. **(d)** Higher magnification images of boxed regions from (c) showing localization patterns from multiplexed AZP labeling. Scale bars = 25 *μ*m. **(e-g)** Quantification of relative Z1 (e), Z7 (f), and Z10 (g) intensity, normalized to polyR binding control signal on a per-cell basis across Eml4-Alk tumors, either in the absence of protease inhibitors (Uninhibited; n = 22 tumors) or in the presence of a broad-spectrum cocktail of protease inhibitors (Inhibited; n = 23 tumors) (mean ± s.d.; two-tailed unpaired t-test, \*\*\*\**P* < 0.0001).

### Delineating enzyme class- and cell type-specific activity with AZPs

Having demonstrated that orthogonal AZPs could be simultaneously multiplexed, we next endeavored to show that they could be used to identify protease families and cell compartments contributing to their cleavage. Due to its prominent *in situ* localization pattern and the significant *in vivo* correlation of PP01 with tumor progression, we nominated Z1 for further investigation and sought to understand the biological processes driving cleavage of this peptide (“S1” for cleavage motif; Table S2). Critically, the specific localization pattern identified from multiplexed AZP staining was conserved with application of Z1 alone to Eml4-Alk lung tissues and was absent in healthy lungs (Fig. 4a). While a free polyR binding control stained Eml4-Alk tumors uniformly, Z1 signal exhibited its prominent spindle-like morphology, indicating that this pattern was not due to heterogeneous charge distribution within the tissue (Fig. 4a). To verify that this localization pattern truly reflected specific protease expression by the labeled cells, rather than nonspecific labeling (i.e., due to non-uniform distribution of charge), we precleaved Z1 *in vitro* with recombinant fibroblast activation protein (FAP; Fig. S5) and compared its tissue labeling to that of intact Z1 activated by tissue-resident enzymes (Fig. S6a-b). While intact Z1 maintained its spindle-like spatial pattern, precleaved Z1 exhibited diffuse, uniform labeling that mirrored that of a free polyR, verifying that probe localization depended on local *in situ* activation (Fig. S6a-b).

**Figure 4:**
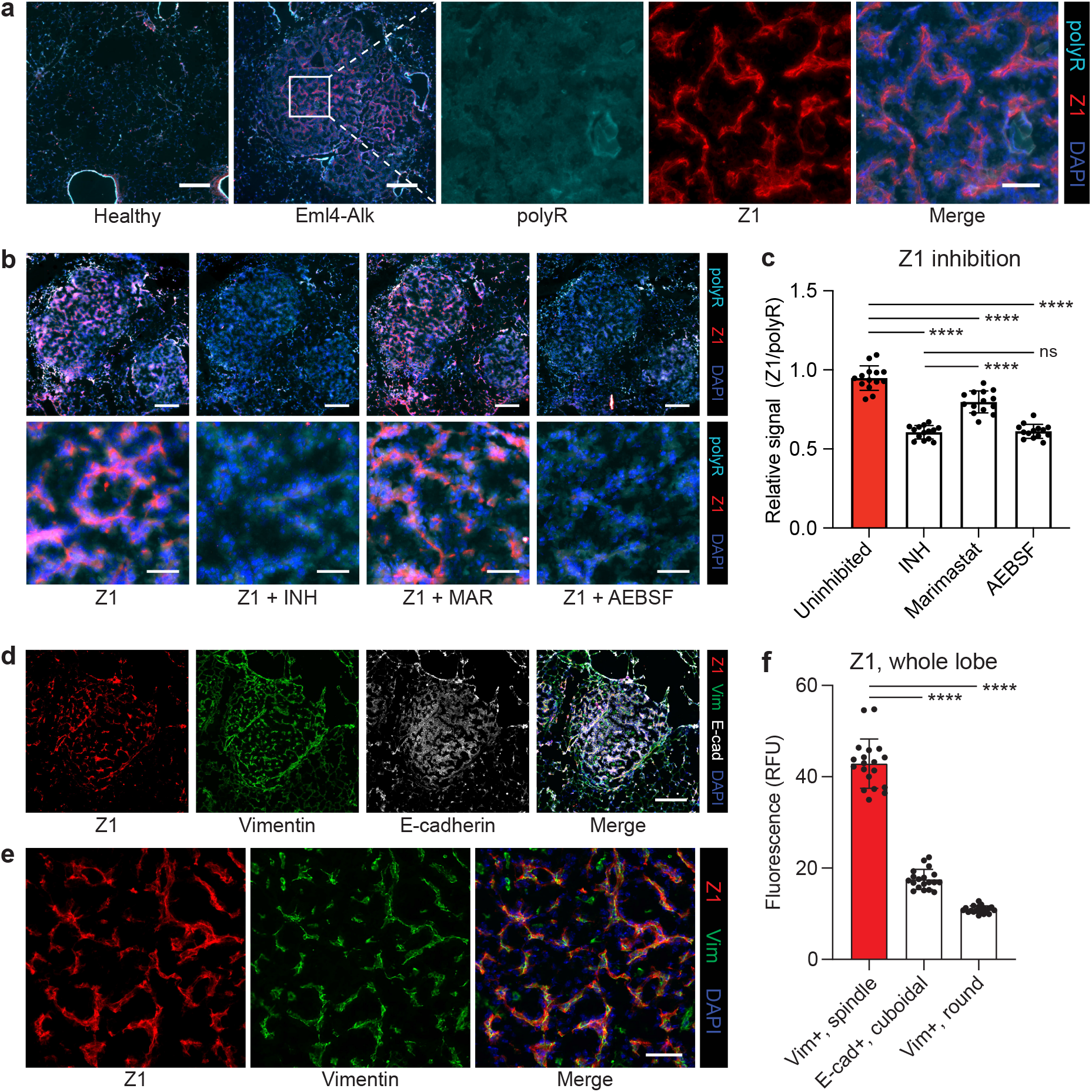
AZPs identify mechanistic class- and cell type-specific protease activity. **(a)** Staining of lung tissue sections from healthy control and Eml4-Alk mice with Z1 (red), polyR (cyan), and DAPI counterstain (blue). Higher magnification images show staining in a representative Eml4-Alk tumor region. Scale bars = 200 *μ*m, 50 *μ*m (lower and higher magnification, respectively). **(b)** Application of Z1 to Eml4-Alk lung tissue sections, either alone (Z1) or in the presence of inhibitors: a broad-spectrum cocktail of protease inhibitors (Z1 + INH), the MMP inhibitor marimastat (Z1 + MAR), or the serine protease inhibitor AEBSF (Z1 + AEBSF). Sections were stained with a polyR binding control (cyan) and counterstained with DAPI (blue). Scale bars = 200 *μ*m (top), 50 *μ*m (bottom). **(c)** Quantification of relative Z1 intensity, normalized to polyR signal, from Eml4-Alk tumors, either in the absence of protease inhibitors (Uninhibited), or in the presence of INH, MAR, or AEBSF (n = 11 tumors; mean ± s.d.; two-tailed unpaired t-test, \*\*\*\**P* < 0.0001, ^ns^*P* = 0.7127). **(d)** Application of Z1 (red) to Eml4-Alk lung tissues sections with co-staining for vimentin (green) and E-cadherin (white). Scale bar = 200 *μ*m. **(e)** Higher-magnification images of a second tumor region, independent of that shown in (d), showing Z1 and vimentin stains. Scale bar = 50 *μ*m. **(f)** Quantification of Z1 staining intensity for per-tumor cell populations, across an entire lung lobe (n = 19 tumors, with intensities averaged across all cells in a tumor; mean ± s.d.; two-tailed paired t-test, \*\*\*\**P* < 0.0001).

To determine class-specific contributions to its cleavage, we applied Z1, whose substrate can be recognized by both matrix metalloproteinases (MMPs) and serine proteases [13, 27, 19], to Eml4-Alk lung tissue sections in the absence of protease inhibitors, with a broad-spectrum cocktail of protease inhibitors, with the MMP inhibitor marimastat, or with the serine protease inhibitor 4-(2-aminoethyl)benzenesulfonyl fluoride hydrochloride (AEBSF; Fig. 4b). As expected, incubation with broad-spectrum protease inhibitors significantly abrogated Z1 labeling (*P* < 0.0001; Fig. 4c). Qualitatively, Z1 signal was largely preserved in sections incubated with marimastat (Fig. 4b). While marimastat did reduce Z1 signal, it remained significantly increased relative to the broad-spectrum inhibitor condition (P < 0.0001; Fig. 4c). In contrast, incubation with AEBSF completely abrogated Z1 tissue labeling to the level of broad-spectrum inhibition (*P* < 0.0001 uninhibited vs. AEBSF, P = 0.7127 broad-spectrum inhibition vs. AEBSF; Fig. 4c), suggesting that serine proteases in Eml4-Alk tumors are primarily responsible for cleaving Z1.

Though Eml4-Alk tumors are adenocarcinomas and thus consist primarily of epithelial cells, the distinct spindle-like labeling pattern of Z1 suggested that Z1 might be cleaved by membrane-bound or secreted proteases expressed by non-epithelial cells of the TME. To this end, we applied Z1 to Eml4-Alk lung tissue sections and co-stained for E-cadherin, an epithelial cell marker, and vimentin, the intermediate filament of non-epithelial, mesenchymal cells (Fig. 4d-e, Fig. S7a-c). We observed strong overlap between Z1 and vimentin (Fig. 4e), with vimentin-positive spindle-like cells exhibiting significantly higher Z1 staining than E-cadherin-positive cells or vimentin-positive rounded cells (likely tumor-associated macrophages) (Fig. 4f; see Methods for details). These results suggest that vimentin-positive, spindle-like cells are responsible for producing the serine proteases that cleave Z1 and, more broadly, demonstrate that AZPs can delineate class-specific and cell type-associated activity patterns.

### Multimodal spatial profiling to functionally query the TME

Next, we explored how enzyme activity profiles measured by our *in vivo* and *in situ* tools correlated with functional and compositional changes within the TME. The distinct spatial pattern of Z1 staining led us to hypothesize that this probe could be labeling cells of the tumor vasculature, rather than cells of immune or other mesenchymal compartments. To this end, we applied Z1 to Eml4-Alk and healthy lungs and co-stained for the endothelial cell marker CD31 (PECAM-1; Fig. S8a). Qualitatively, although both Eml4-Alk and healthy lungs exhibited an abundance of endothelial cells as evidenced by CD31-positivity, Z1 labeling was enriched in Eml4-Alk tumors relative to healthy lungs and tended to colocalize with CD31-positive cells (Fig. 5a). Cell-by-cell quantification of Z1 and CD31 staining intensities across entire lung tissue sections identified a strong positive correlation in Eml4-Alk tissue (*R*^2^ = 0.67; Fig. S8b). Indeed, Z1 staining was significantly increased in CD31-high cells in Eml4-Alk lung tissue sections relative to CD31-high cells in healthy lungs, as well as to CD31-low cells in both Eml4-Alk and healthy tissue (*P* < 0.0001; Fig. 5b), suggesting specific activity associated with the Eml4-Alk tumor endothelium. Furthermore, immunostaining for VE-cadherin, a strictly endothelial-specific adhesion molecule [28], revealed a spindle-like pattern of expression within Eml4-Alk tumors that mimicked Z1 staining (Fig. S9).

**Figure 5:**
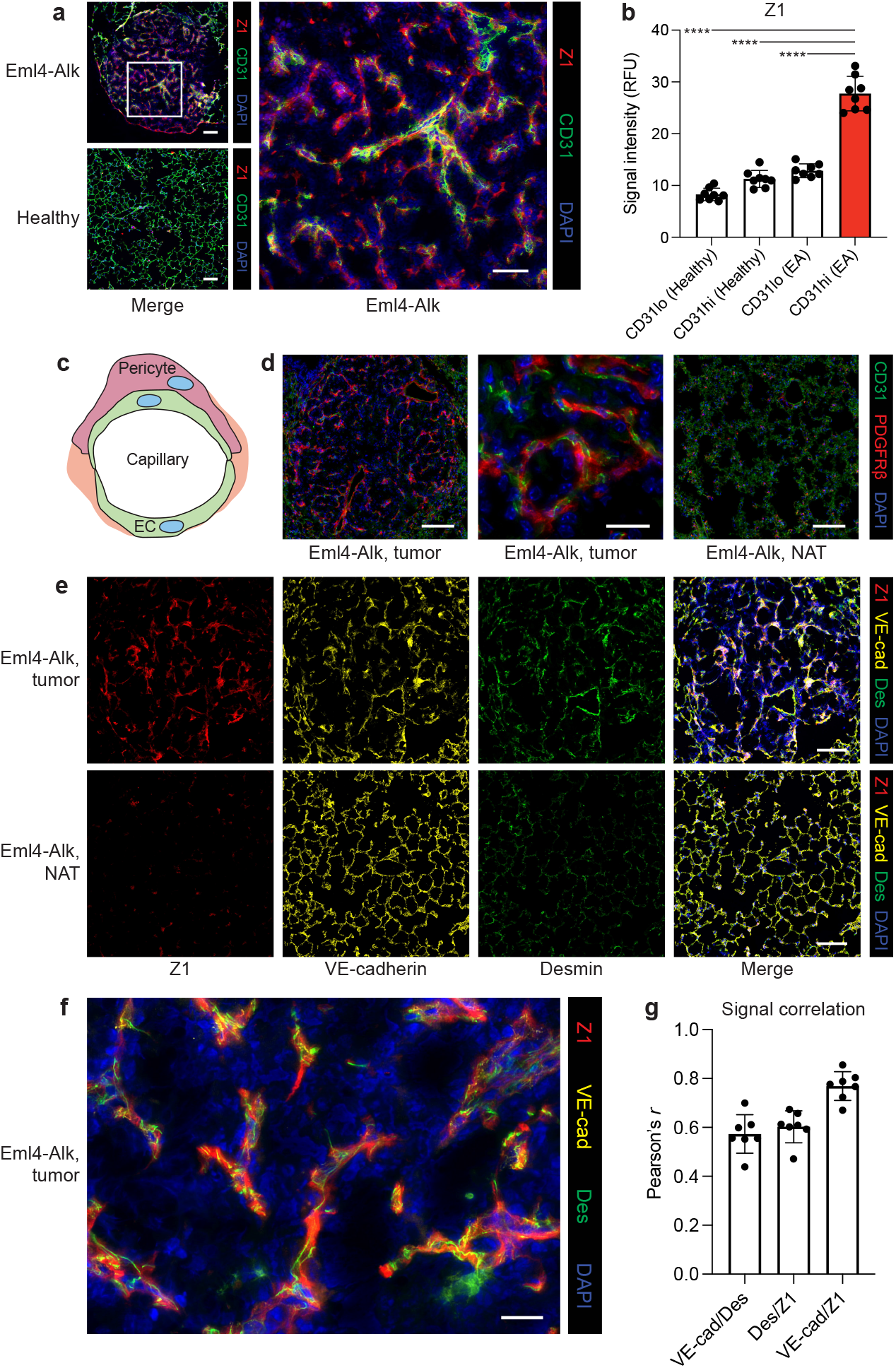
Spatial activity profiling with AZPs enables functional characterization of the tumor microenvironment. **(a)** Application of Z1 (red) to Eml4-Alk and healthy lung tissue sections with co-staining for CD31 (green). Scale bars = 100 *μ*m, 50 *μ*m (lower and higher magnification, respectively). **(b)** Quantification of Z1 staining intensity in CD31-low (CD31lo) and CD31-high (CD31hi) cells (n = 8 regions per condition; mean ± s.d.; one-way ANOVA with Tukey correction for multiple comparisons, ****P < 0.0001). **(c)** Capillary vessels are lined by endothelial cells (EC); pericytes support and wrap around endothelial cells. **(d)** Staining for endothelial cells (CD31; green) and pericytes (PDGFR*β*; red) in formalin-fixed, paraffin-embedded Eml4-Alk lung tissue sections, with images from representative tumor (left, middle) and normal adjacent tissue (NAT; right) regions shown. Scale bar = 100 *μ*m, 20 *μ*m (lower and higher magnification, respectively). **(e)** Application of Z1 (red) to Eml4-Alk lung tissues with co-staining for VE-cadherin (VE-cad; yellow) and desmin (Des; green). Scale bars = 100 *μ*m. **(f)** Higher magnification image of representative Eml4-Alk tumor region showing localization of Z1, VE-cadherin, and desmin. Scale bar = 20 *μ*m. **(g)** Pearson’s correlation of pixel-wise signal intensities for each pairwise combination of Z1, VE-cadherin, and desmin (n = 7 tumors; mean ± s.d.).

In addition to endothelial cells, the vasculature also contains contractile vascular smooth muscle cells that line the vessel walls. Capillaries and microvessels, such as those within the lungs, contain a mural, periendothelial mesenchymal cell population known as pericytes (Fig. 5c), which help regulate vascular function and can be actively recruited into the vasculature during angiogenesis [29, 30, 31]. Eml4-Alk tumors stained positively for *α*-smooth muscle actin (*α*SMA; Fig. S10a), a canonical vascular smooth muscle cell marker that can be expressed by tumor pericytes but is often absent in quiescent pericytes in normal tissues [29, 30]. Indeed, normal adjacent tissue (NAT) showed reduced *α*SMA expression (Fig. S10a). To further corroborate the likely presence of pericytes within the tumor vasculature, we stained Eml4-Alk lungs for CD31 and a second pericyte marker, the muscular intermediate filament desmin [29], and observed desmin-positive cells surrounding CD31-positive endothelial cells within Eml4-Alk tumors but not in NAT (Fig. S10b). Finally, we stained Eml4-Alk lung tissue sections for the pericyte-specific marker PDGFR*β*. The PDGF-B/PDGFR*β* signaling pathway is a key axis of interaction between endothelial cells and pericytes, wherein PDGF-B released from angiogenic endothelial cells binds to PDGFR*β* on the surface of pericytes, facilitating their recruitment [32, 29]. Eml4-Alk tumors stained positively for both CD31 and PDGFR*β*, while NAT from Eml4-Alk lungs did not express PDGFR*β* despite abundant CD31 expression (Fig. 5d, Fig. S10c). Within the tumor vasculature specifically, PDGFR*β*-positive cells wrapped around CD31-positive cells, consistent with the expected localization and function of pericytes (Fig. 5d, Fig. S10c).

To assess its localization with respect to cells of the tumor vasculature, we applied Z1 to Eml4-Alk lung tissue sections with concurrent staining for both the endothelial marker VE-cadherin and the pericyte marker desmin. We observed robust Z1 labeling together with VE-cadherin and desmin expression within Eml4-Alk tumors (Fig. 5e). However, NAT displayed decreased Z1 and desmin staining despite maintaining VE-cadherin positivity. Qualitatively, closer inspection of Z1 labeling within Eml4-Alk tumor regions revealed a close association between all three stains (Fig. 5f). Colocalization analysis demonstrated a correlation between desmin and VE-cadherin staining, consistent with the close proximity of both cell types within capillaries, and additionally showed that both desmin and VE-cadherin correlated with Z1 labeling (Fig. 5g). Together, these results indicate greater abundance of pericytes within the tumor vasculature; suggest that cells of the Eml4-Alk tumor vasculature produce serine proteases that cleave S1; and thus demonstrate that AZPs can be coupled with spatial proteomic approaches to query functional and compositional dysregulation directly within the TME.

### Relating enzyme activity measurements to single-cell transcriptomics for guided biological discovery

After identifying the tumor-specific cellular compartments associated with S1 cleavage, we endeavored to further characterize the phenotypes of the identified vasculature-associated cell populations in order to understand potential mechanisms for their dysregulation. To complement our *in vivo* and *in situ* activity measurements, we performed single-cell RNA sequencing (scRNA-seq) to obtain an unbiased view of the cellular and transcriptomic landscape of Eml4-Alk lungs (Fig. 6a). Graph-based clustering of uniform manifold approximation and projection (UMAP) captured the transcriptomic landscape of Eml4-Alk lungs, where we annotated 8 significant groups of cell types based on expression of previously reported marker genes [33, 34] (Fig. 6b). Given that S1 cleavage *in situ* localized to cells of the tumor vasculature, we defined marker gene modules for both endothelial and pericyte populations and computed their expression scores across all cells in Eml4-Alk lungs (Fig. 6c). The identified population of endothelial cells exhibited robust expression of a module of 28 genes canonically associated with angiogenesis (Fig. S11a-b, Fig. S12). Marker gene analysis additionally revealed a small population of pericytes within the larger stromal cluster (Fig. 6c, Fig. S13a-b).

**Figure 6:**
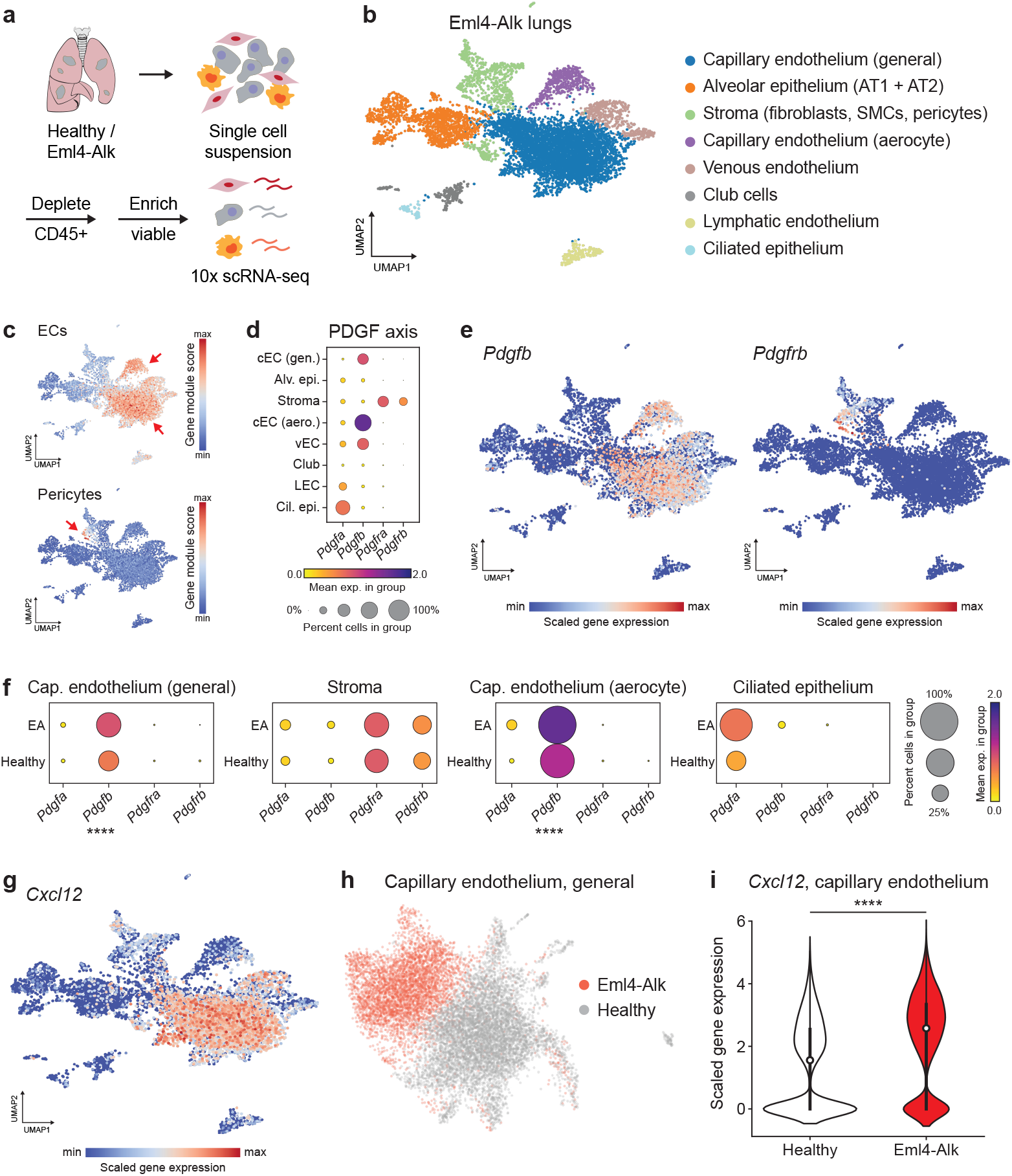
Single-cell transcriptomics for targeted exploration of hypotheses generated by activity profiling. **(a)** Schematic of workflow. **(b)** UMAP of scRNA-seq dataset from Eml4-Alk lungs (pooled sample from n = 3 mice). Cell types were inferred based on expression of canonical marker genes. AT1, alveolar type 1; AT2, alveolar type 2; SMC, smooth muscle cell. **(c)** Feature plots of gene expression module scores for endothelial cell (EC) and pericyte marker genes mapped onto the UMAP of cells from Eml4-Alk lungs. **(d)** Relative expression levels of PDGF and PDGFR genes across cell types in Eml4-Alk lungs. Individual dots represent mean expression values across all cells in a cluster, are colored by expression level, and are sized by the percentage of cells in the cluster expressing that gene. **(e)** Expression levels of *Pdgfb* and *Pdgfrb* against the UMAP of cells from Eml4-Alk lungs. **(f)** Relative expression levels of PDGF and PDGFR genes in cells from Eml4-Alk (EA) and healthy (WT) lungs withinin each of the capillary endothelium (general), stromal, capillary endothelium (aerocyte), and ciliated epithelium populations (Wilcoxon rank-sum test with Benjamini-Hochberg correction, \*\*\*\**P_adj_* < 0.0001). **(g)** Expression levels of *Cxcl12* against the UMAP of cells from Eml4-Alk lungs. **(h)** UMAP of integrated dataset of cells from capillary endothelium (general) populations in Eml4-Alk and healthy lungs (pooled samples of n = 3 mice per condition). **(i)** *Cxcl12* expression in capillary endothelial cells from Eml4-Alk and healthy lungs (Wilcoxon rank-sum test with Benjamini-Hochberg correction, log_2_ FC = 1.453,**** *P_adj_* < 0.0001).

Spatial profiling had indicated the presence of cells positive for each of *α*SMA, desmin, and PDGFR*β* within Eml4-Alk tumors but not within NAT (Fig. S10, Fig. 5c), raising the question of whether pericytes were specifically recruited into the TME. To determine whether pericytes were present in completely normal lungs, we queried scRNA-seq data from healthy mouse lungs for the pericyte marker gene module and identified a small population of cells exhibiting this signature (Fig. S13c-d), in line with pericyte identification reported in previous lung cell atlas studies in human [34] and mouse [33]. Irrespective of the presence of pericytes in completely healthy lungs, PDGF signaling has been shown to facilitate recruitment of pericytes into the tumor vasculature as a means to stabilize vessels and promote the establishment of an angiogenic, reactive TME [35, 36]. We therefore queried expression of both PDGF ligands (*Pdgfa, Pdgfb*) and receptors (*Pdgfra, Pdgfrb*) in the Eml4-Alk scRNA-seq dataset and found that expression of *Pdgfra* and *Pdgfrb* was exclusive to the stromal cluster (Fig. 6d). This analysis also revealed robust and specific expression of *Pdgfb* in endothelial cell populations (Fig. 6d). Visualization via UMAP corroborated that expression levels of *Pdgfb* and *Pdgfrb* matched the distributions of endothelial cell and pericyte populations, respectively (Fig. 6e).

These results raised the possibility of paracrine PDGF signaling between endothelial cells and PDGFR-positive stromal cells in Eml4-Alk tumors. To investigate whether this axis was transcriptionally dysregulated, we conducted an integrative analysis of scRNA-seq data from Eml4-Alk and healthy lungs (Fig. S14a-b). Differential gene expression analysis across this integrated dataset showed that *Pdgfb* was significantly overexpressed in cells from both capillary endothelial cell compartments in the TME relative to healthy lungs (*P_adj_* < 0.0001; Fig. 6f, Table S3). However, expression of the PDGF receptors *Pdgfra* and *Pdgfrb* remained consistent between the total stromal populations from both conditions (Fig. 6f, Table S3).

These observations motivated the hypothesis that altered ligand expression by endothelial cells in the Eml4-Alk TME could be implicated in the association of pericytes to the tumor vasculature. In addition to *Pdgfb*, the chemokine *Cxcl12*, shown to play functional roles in angiogenesis and vascular recruitment of stromal cells [37], was robustly expressed in endothelial cells from Eml4-Alk lungs (Fig. 6g). Endothelial cells from Eml4-Alk and healthy lungs exhibited differential transcriptional landscapes (Fig. 6h), with *Cxcl12* expression significantly increased in endothelial cells from Eml4-Alk lungs relative to those from healthy controls (log_2_ FC = 1.453, *P_adj_* < 0.0001; Fig. 6i, Table S3). Intriguingly, previous reports have shown that overexpression of PDGF-B can increase tumor pericyte content via induction of CXCL12 expression by endothelial cells within the TME [37]. Taken together with our findings from spatial profiling (Fig. 5, Fig. S10), these results raise the possibility that aberrant PDGF and CXCL12 paracrine signaling could potentially help promote increased pericyte coverage of the Eml4-Alk tumor vasculature.

### Activity-based cell sorting for multimodal phenotypic characterization of cancer

Together, these results show that spatial activity profiling can be used to identify the tissue compartment (i.e., the tumor vasculature) associated with a cleavage signature of interest (i.e., of S1), and that complementary omics approaches such as scRNA-seq can be leveraged to follow up on generated hypotheses (i.e., recruitment of pericytes into the tumor vasculature). However, the methods remain decoupled, and thus enzyme activity measurements cannot be directly linked to other measurement modalities at the cellular level. We therefore sought to establish a method to isolate individual cells purely on the basis of an enzymatic activity signature. We hypothesized that AZPs containing fluorophore-quencher pairs could function as enzyme-activatable cellular tags *in vivo* to label cells with membrane-bound or proximal protease activity, such that tagged cells could be subsequently sorted via flow cytometry (Fig. 7a). In this design, following systemic administration, degradation of the protease-cleavable linker activates fluorescence and liberates the fluorophore-tagged polyR such that it can bind and tag nearby cells, functioning analogously to a cell penetrating peptide [38, 26]. Thus, we reasoned that probe labeling after proteolytic activation (e.g., by cell-surface or proximal proteases) would facilitate isolation of tagged cells via fluorescence-activated cell sorting (FACS) and enable subsequent downstream phenotypic characterization (Fig. 7a).

**Figure 7:**
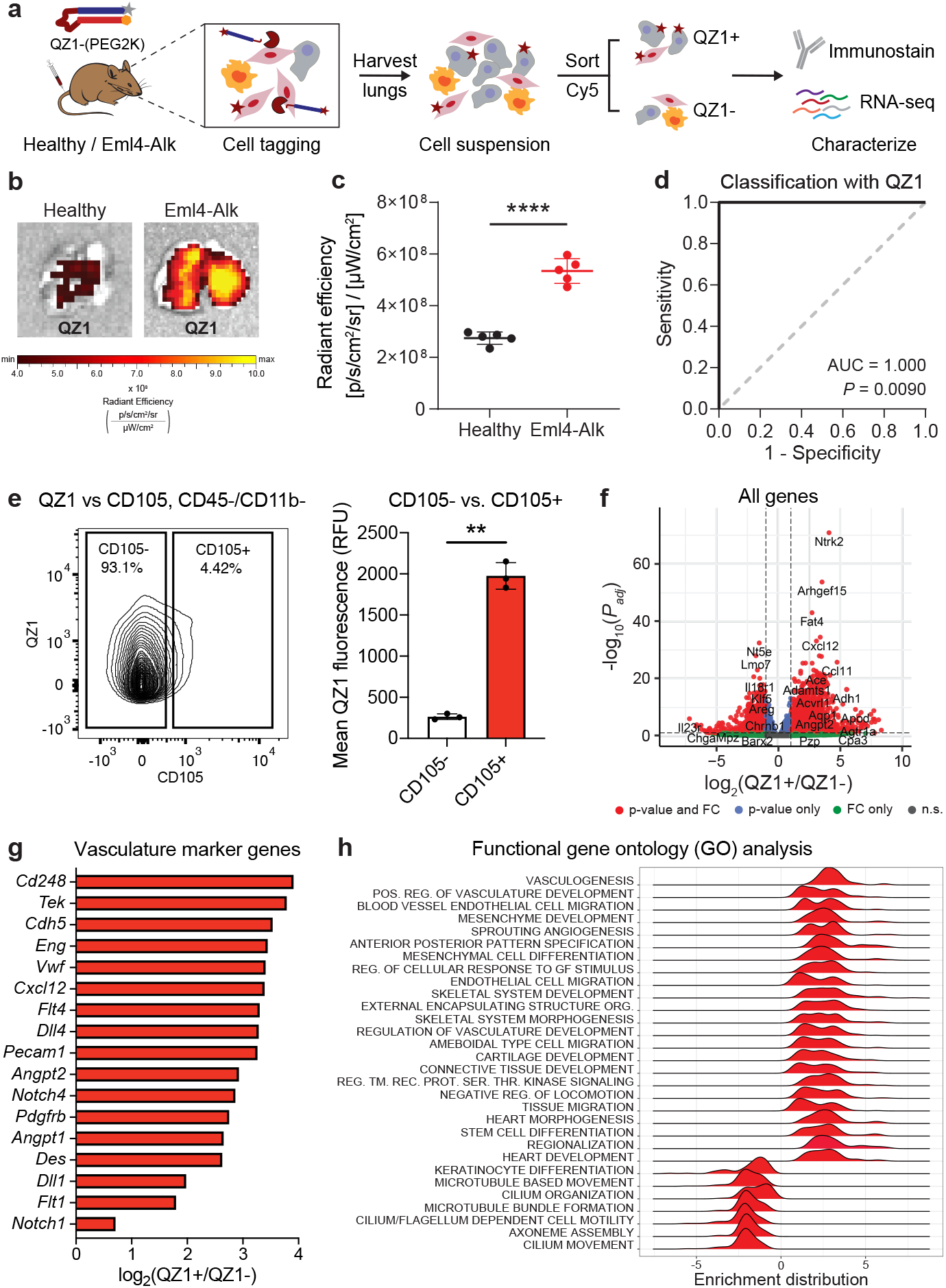
Activity-based cell sorting enables multimodal phenotypic characterization of cancer. **(a)** The quenched probe QZ1-(PEG2K), consisting of a Cy5-tagged polyR (gray star + blue rectangle) and quencher-tagged polyE (orange hexagon + red rectangle), was administered intravenously into age-matched healthy and Eml4-Alk mice. Lungs were excised, imaged, and dissociated into single cell suspension. Cells were sorted on Cy5 fluorescence and then characterized via immunostaining and bulk RNA-seq. **(b)** Images of representative lungs from healthy and Eml4-Alk mice 2 hours after QZ1-(PEG2K) administration. **(c)** Quantification of epifluorescence radiant efficiency from experiment in (b) (n = 5 mice per group; mean ± s.d.; two-tailed unpaired t-test, \*\*\*\**P* < 0.0001). **(d)** ROC curve showing performance of QZ1-(PEG2K) signal in discriminating healthy from Eml4-Alk lung explants (AUC = 1.000, 95% confidence interval 1.000–1.000; P = 0.0090 from random classifier shown in dashed line). **(e)** Flow cytometry plot (left) and quantification (right) of QZ1 fluorescence intensity in CD45-, CD11b-cells from Eml4-Alk lungs, gated by endoglin expression (CD105; n = 3 biological replicates; mean ± s.d.; two-tailed paired t-test, **P = 0.00285). **(f)** Differential expression analysis of RNA-seq data from sorted QZ1+ and QZ1-cells. Each point represents one gene and is colored according to the significance level for that gene. **(g)** Identification of vasculature marker genes significantly overexpressed (*P_adj_* < 0.05) in the QZ1+ population. **(h)** Enrichment distributions for gene ontology modules up- and downregulated in the QZ1+ population relative to the QZ1-population.

We applied this activity-based cell sorting assay to directly isolate and then phenotypically characterize the Eml4-Alk cell compartment associated with S1 cleavage (Fig. 7a). We designed a fluorescent quenched probe, QZ1, that incorporated S1 as a protease-cleavable linker. Cy5-labeled QZ1 was PEGylated to improve stability and drive tissue accumulation [38, 26], and administered intravenously into age-matched Eml4-Alk and healthy mice. Eml4-Alk mice were assessed at 12 weeks post tumor induction, at which point lungs contain multiple lung adenocarcinoma lesions [23]. Two hours post injection, significantly increased Cy5 fluorescence was found in explanted Eml4-Alk lungs relative to healthy lungs (P < 0.0001; Fig. 7b-c), enabling perfect discrimination of Eml4-Alk and healthy mice (AUC = 1.000; Fig. 7d).

Following imaging, single-cell suspensions were prepared from dissociated Eml4-Alk lungs, and flow cytometry was used to sort all live, non-hematopoetic nucleated cells by QZ1 signal, validating the feasibility of the activity-based cell sorting method (Fig. 7a, Fig. S15). Immunostaining concurrent to the activity-based sort revealed significantly increased QZ1 signal in cells positive for the endothelial cell marker endoglin (CD105; P = 0.00285; Fig. 7e). Increased Cy5 signal was also observed in cells positive for CD44 (P = 0.0120; Fig. S16a), which can help regulate division and angiogenesis in endothelial cells [39], and for Ly-6A/E (P = 0.000374; Fig. S16b), a marker of hematopoietic cells [40] that can also be expressed by pulmonary endothelium [41].

Bulk RNA-seq on sorted QZ1-positive (QZ1+) and QZ1-negative (QZ1-) populations was used to characterize gene expression differences between the two compartments (Fig. 7f, Fig. S17). Several canonical markers of endothelial cells (*Cdh5, Eng, Vwf, Pecam1*), pericytes (*Cd248, Pdgfrb, Des*), as well as vascularization and angiogenesis were among the most upregulated genes in the QZ1+ population (Fig. 7g). Gene set enrichment analysis corroborated that the dominant cell types associated with QZ1 positivity were endothelial and mesenchymal cell types, including pericytes, while gene sets associated with epithelial cells were significantly downregulated (Fig. S18a). Critically, *Cxcl12* and *Pdgfrb* were overexpressed in the QZ1+ compartment, as were markers of additional angiogenesis signaling axes, including the VEGF (*Flt1/4*), Notch (*Notch1/4, Dll1/4*), and Tie (*Tek, Angpt1/2*) pathways (Fig. 7g). Indeed, the QZ1+ compartment was significantly enriched for functional modules associated with vasculogenesis, vascular development, endothelial cell migration, and mesenchymal recruitment (Fig. 7h, Fig. S18b). More generally, these results validate an activity-based cell sorting assay that can be directly coupled with protein- and transcript-level measurements for multimodal phenotypic characterization.

## Discussion

### Summary

Our results establish a paradigm for profiling enzyme activity across multiple scales–at the organism, tissue, and cellular levels–and demonstrate the utility of our methods for noninvasive monitoring and functional characterization of cancer (Fig. 1). We first demonstrated that multiplexed panels of protease-responsive nanosensors can quantitatively track disease dynamics *in vivo* to yield activity-based biomarkers of tumor progression and targeted therapy response. Next, we turned to the spatial setting and directly translated substrates nominated from *in vivo* profiling into *in situ* protease activity probes (AZPs). These methods enabled identification of a tumor-specific serine protease activity signal that increased with tumor progression, rapidly decayed after therapy, and localized specifically to the pericyte-invested tumor vasculature. We complemented our activity measurements with single-cell transcriptomic analysis, which identified overexpression of paracrine signaling factors in endothelial cells from tumor-bearing lungs and suggested a possible mechanism for endothelial cell-pericyte crosstalk within the TME. Finally, we designed a high-throughput method to isolate cells based on their protease activity and leveraged it to discover a population of proteolytically active, vasculature-associated cells harboring pro-angiogenic transcriptional programs. Together, these methods revealed that the functionally aberrant tumor vasculature rapidly responds to tumor cell-targeted inhibition of oncogenic signaling and demonstrated that protease activity serves as an informative proxy for this process. By designing our methods to uniformly rely on a single measurement mechanism (i.e., proteolytic cleavage of a peptide substrate), we establish methodology for enzyme activity profiling across biological scales, with both temporal and spatial resolution.

### Next-generation methods for profiling enzyme activity in cancer

Our work establishes the first multiplexed *in situ* activity assay that enables direct on-tissue comparison of spatial localization patterns of distinct proteases. Further expansions of AZP multiplexing capacity, either through epitope tagging or DNA barcoding, could enable high-throughput screens on murine or human tissue to discover enzyme activity sensors with desired properties, such as colocalization with specific cell types. We also demonstrate that AZPs can be used to delineate protease class-specific activity signatures through targeted inhibitor ablations *in situ*. Given that our results indicate that serine proteases cleave S1 in the Eml4-Alk model (Fig. 4c), additional molecular profiling will be necessary to identify which protease(s) are responsible. The angiogenesis-associated proteases *Plat* (tPA) and *Dpp4* (DPP4), as well as the membrane serine protease *Fap* (FAP or seprase), which can be selectively expressed by tumor pericytes [42, 43], were amongst the serine proteases overexpressed in the QZ1+ population (Fig. S17). *In vitro* screening, targeted small molecule inhibition, or specific knockout of individual enzyme targets could help identify the serine protease(s) that cleave S1 in the Eml4-Alk model. Parallel functional studies may help determine the specific contributions of candidate serine proteases to vascular remodeling and angiogenesis in ALK^+^ lung cancer.

Perhaps most importantly, we establish AZPs as an activity-based cellular tag for sorting individual cells based on endogenous enzyme activity. Administration of AZPs *in vivo*, followed by tissue dissociation and FACS, enabled isolation of cells exhibiting a specific pattern of protease activity. By coupling this assay to immunostaining and RNA-seq, we demonstrate that activity-based cell sorting can enable multimodal characterization of cancer across the activity, protein, and gene expression levels. Probes similar in concept to AZPs could extend activity-based cell sorting to other classes of enzymes. In addition, integrating activity-based cell sorting with large-scale omics measurements and machine learning could inspire a new class of single-cell multiomics that ends at the level of actuated biological function. We envision that the ability to sort cells by enzymatic activity could yield transformative insights into enzymatic dysregulation in disease; enable multimodal approaches to comprehensively characterize biological systems; and inform new diagnostic and therapeutic interventions.

### Multimodal profiling to explore cancer biology

Our results demonstrate that protein activity directly complements measurements of protein and transcript abundance, and that this multimodal profiling enables discovery-based functional characterization of the TME. By applying our activity-based profiling methods to the Eml4-Alk model of NSCLC, we discover aberrant serine protease activity that is specific to the tumor vasculature and rapidly responds to inhibition of an adjacent cancer-cell specific pathway. Through a combination of spatial profiling and scRNA-seq analysis, we found evidence suggestive of increased pericyte coverage within the Eml4-Alk tumor vasculature, potentially mediated by altered paracrine signaling via PDGF and CXCL12. Though mechanistic experiments will be necessary to ascertain whether pericytes are actively recruited into the TME, our findings raise the possibility that S1 cleavage, which is elevated within Eml4-Alk tumors and localizes specifically to the vasculature, could be due to the coordinated action of intratumoral pericytes and endothelial cells associated with neoangiogenic vessels. Our finding that the functionally aberrant tumor vasculature rapidly responds to targeted therapy motivates exploration of whether anti-angiogenic drugs, which have been clinically approved in combination with cytotoxic chemotherapy or immunotherapy [44, 45, 46], could have additive benefits when combined with molecularly targeted therapeutics like alectinib. Our study in the Eml4-Alk model serves as an example for how our activity profiling methods can be leveraged to advance understanding of the complex crosstalk between cancer and non-cancer cells. We envision that integrating these activity profiling techniques with other data modalities will enable a more complete understanding of tumor biology.

### Opportunities and applications in precision cancer medicine

Finally, the activity-based profiling methods presented here could have significant utility in precision medicine applications. Precision cancer medicine requires granular information that cannot be accessed by traditional noninvasive imaging approaches, necessitating serial biopsies that carry significant risks and sample only a small fraction of the disease site. The ability to gain high-dimensional biological insight into a disease state with a completely noninvasive test would present a paradigm shift towards functional precision medicine [47]. Here, we establish the ability of noninvasive, multiplexed protease activity nanosensors to query the function and activity of specific intratumoral cell subsets over the course of tumor progression and in response to therapy. Given the modularity of this approach, high-throughput screening [48, 49, 50] and generative machine learning [51] methods could optimize sensors to target orthogonal axes of cancer biology. For instance, sensors that detect angiogenesis could be administered in combination with probe sets that read out immune invasion or metastasis risk. As a complement to this noninvasive test, a targeted panel of *in situ* AZPs could be used to molecularly profile individual patient biopsies for indication of signaling pathways or processes active in a patient’s specific tumor. This information would empower patients and physicians with real-time, high-quality information to personalize treatment decisions, such as rapid prediction of immunotherapy efficacy, surveillance for recurrence after targeted therapy, or discrimination of aggressive versus indolent disease.

### Conclusion

In summary, we have developed a integrated suite of enzyme activity-profiling methods that form a direct link between noninvasive enzyme sensors, high-resolution spatial profiling, and high-throughput, single-cell analytical methods like flow cytometry and RNA-seq. The modular methods described here can be readily generalized to other cancer types and hold promise for both fundamental biological investigation and translational research. We envision that these methods for profiling enzyme activity will enable a more comprehensive assessment of tumor biology and facilitate functional characterization of cancer for medical and discovery applications alike.

## Materials and Methods

### Eml4-Alk mouse model of non-small-cell lung cancer

All animal studies were approved by the Massachusetts Institute of Technology (MIT) committee on animal care (protocol 0420-023-23) and were conducted in compliance with institutional and national policies. Reporting was in compliance with Animal Research: Reporting *In Vivo* Experiments (ARRIVE) guidelines. Tumors were initiated in female C57BL/6 mice between 6 and 10 weeks old by intratracheal administration of 50 *μ*L adenovirus expressing the Ad-EA vector (Viraquest, 1.5 * 10^8^ PFU in Opti-MEM with 10 mM CaCl_2_), as described previously [23]. Healthy control cohorts consisted of age- and sex-matched mice that did not undergo intratracheal administration of Ad-EA adenovirus.

### Alectinib treatment

Eml4-Alk mice were randomized to receive either control vehicle or alectinib (MedChemExpress), at 20 mg/kg prepared directly in drug vehicle, daily by oral gavage for 14 consecutive days. Drug vehicle consisted of: 10% (v/v) dimethylsulfoxide (DMSO; Sigma Aldrich), 10% (v/v) Cremophor EL (Sigma Aldrich), 15% (v/v) poly(ethylene glycol)-400 (PEG400; Sigma Aldrich), 15% (w/v) (2-Hydroxypropyl)-*β*-cyclodextrin; Sigma Aldrich). Mice were monitored daily for weight loss and clinical signs. Investigators were not blind with respect to treatment.

### *In vivo* characterization of activity-based nanosensors

All activity-based nanosensor experiments were performed in accordance with institutional guidelines. Tumor-bearing mice and age-matched controls were administered activity-based nanosensor constructs via intratracheal intubation at 3.5, 5, 5.5, 6, and 7 weeks after tumor induction, with treatment of vehicle control or alectinib beginning at 5 weeks after tumor induction in Eml4-Alk mice and continuing for 2 weeks. Nanosensors for urinary experiments were synthesized by CPC Scientific. The urinary reporter glutamate-fibrinopeptide B (Glu-Fib) was mass barcoded for detection by mass spectrometry. Sequences are provided in Table S1. Nanosensors were dosed (50 *μ*L total volume, 20 *μ*M each nanosensor) in mannitol buffer (0.28 M mannitol, 5 mM sodium phosphate monobasic, 15 mM sodium phosphate dibasic, pH 7.0-7.5) by intratracheal intubation. Anesthesia was induced by isoflurane inhalation, and mice were monitored during recovery. For intratracheal instillation, a volume of 50 *μ*L was administered by passive inhalation following intratracheal intubation with a 22G flexible plastic catheter (Exel). Intratracheal instillation was immediately followed by a subcutaneous injection of PBS (200 *μ*L) to increase urine production. Bladders were voided 60 minutes after nanosensor administration, and all urine produced 60-120 min after administration was collected using custom tubes in which the animals rest upon 96-well plates that capture urine. Urine was pooled and frozen at °80°C until analysis by LC-MS/MS.

### LC-MS/MS reporter quantification

LC-MS/MS was performed by Syneos Health using a Sciex 6500 triple quadrupole instrument. Briefly, urine samples were treated with ultraviolet irradiation to photocleave the 3-Amino-3-(2-nitro-phenyl)propionic Acid (ANP) linker and liberate the Glu-Fib reporter from residual peptide fragments. Samples were extracted by solid-phase extraction and analyzed by multiple reaction monitoring by LC-MSMS to quantify concentration of each Glu-Fib mass variant. Analyte quantities were normalized to a spiked-in internal standard and concentrations were calculated from a standard curve using peak area ratio (PAR) to the internal standard. Mean scaling was performed on PAR values to account for mouse-to-mouse differences in activity-based nanosensor inhalation efficiency and urine concentration.

### Statistical and machine learning analysis of urinary reporter data

For all urine experiments, PAR values were normalized to nanosensor stock concentrations and then mean scaled across all reporters in a given urine sample prior to further statistical analysis. To identify differential urinary reporters, reporters were subjected to unpaired two tailed t-test followed by correction for multiple hypotheses using the Holm-Sidak method in GraphPad Prism 9.0. *P_adj_* < 0.05 was considered significant. PCA was performed on mean scaled PAR values and implemented in MATLAB R2019b (Mathworks). For treatment response classification based on urinary activity-based nanosensor signatures, randomly assigned sets of paired data samples consisting of features (the mean scaled PAR values) and labels (for example, EA, Alectinib) were used to train random forest (36) classifiers implemented with the TreeBagger class in MATLAB R2019b. Estimates of out-of-bag error were used for cross-validation, and trained classifiers were tested on randomly assigned, held-out, independent test cohorts. Ten independent train-test trials were run for each classification problem, and classification performance was evaluated with ROC statistics calculated in MATLAB. Classifier performance was reported as the mean accuracy and AUC across the ten independent trials.

### AZP peptide synthesis and sequences

All AZPs were synthesized by CPC Scientific (Sunnyvale, CA) and reconstituted in dimethylformamide (DMF) unless otherwise specified. AZP sequences are provided in Table S2.

### *In situ* zymography with activatable zymography probes

Mice were euthanized by isoflurane overdose. Lungs were then filled with undiluted optimal-cutting-temperature (OCT) compound through catheterization of the trachea; the trachea was subsequently clamped; and lungs were extracted. Individual lobes were dissected and then immediately embedded and frozen in optimal-cutting-temperature (OCT) compound (Sakura). Cryosectioning was performed at the Koch Institute Histology Core. Prior to staining, slides were air dried, fixed in ice-cold acetone for 10 minutes, and then air dried. After hydration in PBS (3×5 minutes), tissue sections were blocked in protease assay buffer (50 mM Tris, 300 mM NaCl, 10 mM CaCl_2_, 2 mM ZnCl_2_, 0.02% (v/v) Brij-35, 1% (w/v) BSA, pH 7.5) for 30 minutes at room temperature. Blocking buffer was aspirated, and solution containing fluorescently labeled AZPs (1 *μ*M each AZP) and a free poly-arginine control (polyR, 0.1 *μ*M) diluted in the protease assay buffer was applied. Slides were incubated in a humidified chamber at 37° C for 4 hours. For inhibited controls, 400 *μ*M AEBSF (Sigma Aldrich), 1 mM marimastat (Sigma Aldrich), or protease inhibitor cocktail (P8340, Sigma Aldrich) spiked with AEBSF and marimastat was added to the buffer at both the blocking and cleavage assay steps. For uninhibited conditions, dimethyl sulfoxide (DMSO) was added to the assay buffer to a final concentration of 3% (v/v). For co-staining experiments, primary antibodies (E-cadherin, AF748, R&D Systems, 4 *μ*g/mL; vimentin, ab92547, Abcam, 0.5 *μ*g/mL; CD31, AF3628, R&D Systems, 10 *μ*g/mL; desmin, ab227651, Abcam, 1.32 *μ*g/mL) were included in the AZP solution. Following AZP incubation, slides were washed in PBS (3×5 minutes), stained with Hoechst (5 *μ*g/mL, Invitrogen) and the appropriate secondary antibody if relevant (Invitrogen), washed in PBS (3×5 minutes), and mounted with ProLong Diamond Antifade Mountant (Invitrogen). Slides were scanned on a Pannoramic 250 Flash III whole slide scanner (3DHistech).

### AZP precleavage characterization

The Z1 AZP (10 *μ*mol/L) was incubated with recombinant fibroblast activation protein (FAP) in FAP assay buffer (50 mM Tris, 1 M NaCl, pH 7.5) overnight at 37° C to run the cleavage reaction to completion. After precleavage with recombinant FAP, the AZP solution was diluted to a final peptide concentration of 0.1 *μ*M in protease assay buffer. Cognate intact Z1 AZP (1 *μ*mol/L) and precleaved Z1 AZP, each with a free polyR control (0.1 *μ*M), were applied to fresh-frozen Eml4-Alk lung tissue sections (slide preparation described above) and incubated at 37° C for 4 hours. After AZP incubation, slides were washed, stained with Hoechst, mounted, and scanned.

### Immunohistochemistry and immunofluorescence staining

Lungs were excised and either embedded in OCT, as previously described, or fixed in 10% (v/v) formalin and embedded in paraffin. Prior to staining, slides with formalin-fixed, paraffin-embedded sections were subject to deparaffinization and antigen retrieval. Prior to staining, slides with fresh-frozen sections were air dried, fixed in ice-cold acetone for 10 minutes, air dried, and re-hydrated in PBS. Sections were stained with IgG isotype controls (ThermoFisher) and primary antibodies against vimentin (ab92547, Abcam, 1.0 *μ*g/mL), E-cadherin (AF748, R&D Systems, 4.0 *μ*g/mL), *α*-SMA (ab124964, Abcam, 1.5 *μ*g/mL), CD31 (AF3628, R&D Systems, 10 *μ*g/mL), VE-cadherin (36-1900, Invitrogen, 10 *μ*g/mL), PDGFR*β* (3169, Cell Signaling, 1: 100), and desmin (ab227651, Abcam, 1.32 *μ*g/mL), as appropriate. For immunohistochemistry, slides were incubated with the appropriate secondary antibody conjugated to horseradish peroxidase (HRP). For immunofluorescence, slides were washed in PBS, incubated with the appropriate secondary antibody and Hoechst, and washed in PBS. Slides were scanned as previously described.

### Quantification of AZP and immunofluorescence staining

AZP and immunofluorescence staining was quantified in QuPath 0.2.3[52] and in ImageJ. To perform cell-by-cell analysis, cell segmentation was performed using automated cell detection on the DAPI (nuclear) channel. For quantification of activity inhibition, AZP staining was calculated as a fold change of the mean nuclear AZP signal over the mean nuclear polyR signal. All nuclei within an individual tumor were averaged across that given tumor. Nuclei with a polyR intensity of less than 3 were excluded from analysis. For quantification of AZP intensity based on cell morphology and marker expression, cells were annotated as “vimentin-positive, spindle” if they were spindle-shaped and expressed vimentin; “E-cadherin-positive, cuboidal” if they were cuboidal-shaped and expressed E-cadherin; “vimentin-positive, round” if they were rounded and expressed vimentin. A random forest classifier was trained on all annotated cells (at least 20 cells per class) using multiple cellular features, including nuclear area and eccentricity, and mean cellular fluorescence intensity across all channels. The trained classifier was then applied to all cells across all tumors in the tissue section, and mean cellular fluorescence intensity was quantified. To assess relationship between Z1 and CD31, cell segmentation was performed as described above and correlation was assessed between mean cellular Cy5 (Z1) intensity and mean cellular FITC (CD31) intensity. Density plots were generated using the dscatter function in MATLAB R2019b. For quantification of co-localization, JACoP (Just Another Co-localization Plug-in) [53] was used to determine pixel intensity-based correlations. Tumors were selected as regions of interest, and thresholds were chosen automatically using the Costes’ method. Co-localization was assessed via the pairwise correlation of pixel intensities within each tumor region of interest.

### *In vivo* administration of QZ1

QZ1 (Table S2) was reconstituted to 1 mg/mL in water, then reacted with mPEG-Maleimide, MW 2000 g/mol (Laysan Bio), for PEG coupling via maleimide-thiol chemistry. After completion of the reaction, the final compound was purified using HPLC. All reactions were monitored using HPLC connected with mass spectrometry. Characterization of the final compound, QZ1-(PEG2K), using HPLC and MALDI-MS indicated that products were obtained with more than 90% purity and at the expected molecular weight. Eml4-Alk mice (11–12 weeks post tumor induction) and age-matched C57BL/6 healthy controls (Jackson Labs; 18–22 weeks) were anesthetized using isoflurane inhalation (Zoetis). QZ1-(PEG2K) (4.5 nmoles in 0.9% NaCl) was administered intravenously via tail vein injection. Two hours after probe injection, mice were imaged on an *in vivo* imaging system (IVIS, PerkinElmer) by exciting Cy5 at 640 nm and measuring emission at 680 nm. Mice were subsequently euthanized by isoflurane overdose followed by cervical dislocation. Lungs were dissected and explanted for imaging via IVIS. Fluorescence signal intensity was quantified using the Living Image software (PerkinElmer).

### Preparation of single-cell suspensions

Eml4-Alk mice (10–12 weeks post tumor induction) and age-matched C57BL/6 healthy controls (Jackson Labs; 18–22 weeks) were euthanized by isoflurane overdose, and lungs were excised and separated into lobes. For tumor-bearing lungs, tumors were separated from healthy tissue using a dissecting microscope. Tissue was minced, treated with digestion buffer (Hank’s Balanced Salt Solution + 2% heat-inactivated FBS, supplemented with DNase and collagenase), and incubated at 37C for 30 minutes with rotation. Samples were filtered using a 70 *μ*m filter and diluted with RPMI-1640 + 2% heat-inactivated FBS. Cell suspension was centrifuged at 1800 rpm for 5 minutes and the pellet was resuspended in ACK lysis buffer for 2 minutes, followed by quenching with FACS buffer (PBS + 2% heat-inactivated FBS). Cell suspension was centrifuged and supernatant was discarded. For single cell RNA-Seq, CD45 cell depletion and viability enrichment was performed according to manufacturer’s instructions (StemCell Technologies).

### Activity-based cell sorting

Single cell lung suspensions from Eml4-Alk mice administered QZ1 were stained with the following antibodies (clone, fluorophore, dilution): CD44 (IM7, BV605, 1:200), CD105 (MJ7/18, BV786, 1:200), Ly6-A/E (D7, PE, 1:200), CD11b (M1/70, APC-Cy7, 1:200), CD45 (30-F11, AF488, 1:400), and EpCAM (G8.8, PE-Cy7, 1:200). Cells were stained for 20 minutes, and DAPI (1:10,000) was added immediately prior to sort. FACS sorting was performed on a FACSAria II (BD). Flow cytometry data was analyzed by the FlowJo software (Treestar). The sort strategy is shown in Fig. S15. At least 100,000 cells from each of the QZ1+ and QZ1-compartments were collected into RPMI-1640 + 2% heat-inactivated FBS and pelleted via centrifugation at 1800 rpm for 5 minutes. Cell pellets were lysed in Trizol (ThermoFisher), and RNA was extracted using RNEasy Mini Kits (Qiagen). Bulk RNA sequencing was performed by the MIT BioMicro Center. Libraries were prepared using the Clontech SMARTer Stranded Total RNAseq Kit (Clontech), precleaned, and sequenced using an Illumina NextSeq500 on an Illumina NextSeq flow cell. Feature counting was performed on BAM files using the Rsubread package. Differential expression analysis on QZ1+ vs QZ1-cells was performed using the DESeq2 package in R. GSEA was performed using GenePattern [54], and results were visualized using the clusterProfiler R package.

### Analysis of Eml4-Alk bulk RNA-seq dataset

Differential expression analysis over the entire transcriptome was performed on a bulk RNA-seq dataset from the Eml4-Alk mouse model of NSCLC, reported by Li et al. [24], using the DESeq2 package in R. The gene list was subsequently filtered to protease genes for visualization.

### Single cell RNA sequencing (scRNA-seq)

Single cells were processed using the 10X Genomics Single Cell 3’ platform using the Chromium Single Cell 3’ Library & Gel Bead Kit V2 kit (10X Genomics), per manufacturer’s protocol. Briefly, approximately 10,000 cells were loaded onto each channel and partitioned into Gel Beads in Emulsion (GEMs) in the 10x Chromium instrument. Following lysis of the captured cells, the released RNA was barcoded through reverse transcription in individual GEMs, and complementary DNA was generated and amplified. Libraries were constructed using a Single Cell 3’ Library and Gel Bead kit. The libraries were sequenced using an Illumina NovaSeq6000 sequencer on an Illumina NovaSeq SP flow cell. scRNA-seq was performed by the MIT BioMicro Center.

### Single cell RNA-seq data analysis

Raw gene expression matrices were generated for each sample by the Cell Ranger (v.3.0.2) Pipeline coupled with mouse reference version GRCm38. The output filtered gene expression matrices were analyzed by Python software (v.3.9.0) with the scanpy package (v.1.7.2) [55]. Genes expressed in at least three cells in the data and cells with > 200 genes detected were selected for further analyses. Low quality cells were removed based on number of total counts and percentage of mitochondrial genes expressed. After removal of low quality cells, gene expression matrices were normalized. The dataset was additionally filtered to remove cells expressing *Ptprc* (CD45). Features with high cell-to-cell variation were calculated. To reduce dimensionality, principal component analysis was conducted with default parameters on normalized and scaled data. Following uniform manifold approximation and projection (UMAP) for dimensionality reduction, cells were clustered in the UMAP embedding space using the Louvain algorithm with resolution 0.25, and cell types were annotated based on expression of known lung cell type marker genes curated from the literature. All analyses and visualizations were implemented in Python with support from scanpy [55].

### Statistical analysis

Differential gene expression analysis for bulk RNA-seq data was performed in R. PCA and machine learning classification of activity-based nanosensor data was performed in MATLAB R2019B. scRNA-seq data analysis was performed in Python (v.3.9.0) using the scanpy (v.1.7.2) package. All remaining statistical analyses were conducted in Prism 9.0 (GraphPad). Sample sizes, statistical tests, and p-values are specified in figure legends.

## Data availability

All data is available upon request to.

## Acknowledgements

The authors thank Dr. C. Chen (Boston University, Boston, MA), Dr. I. Dagogo-Jack (Massachusetts General Hospital, Boston, MA), Dr. V. Gocheva (MIT, Cambridge, MA), Dr. R. Hynes (MIT, Cambridge, MA), and Dr. L. Sequist (Massachusetts General Hospital, Boston, MA) for valuable discussion, feedback, and insight; M. Anahtar, C. Martin-Alonso, and E. Sanders (MIT, Cambridge, MA) for technical assistance; A. Mancino (Syneos Health) for performing mass spectrometry; Dr. S. Levine of the MIT BioMicro Center (MIT, Cambridge, MA); K. Cormier of the Koch Institute Hope Babette Tang Histology Core (MIT, Cambridge, MA); and the MIT BioMicro Center and the Koch Institute Swanson Biotechnology Center, specifically the Histology core and the Preclinical Modeling, Imaging, and Testing core. This study was supported, in part, by a Koch Institute Support Grant P30-CA14051 from the National Cancer Institute, a Core Center Grant P30-ES002109 from the National Institute of Environmental Health Sciences, the Virginia and D.K. Ludwig Fund for Cancer Research, the Koch Institute Frontier Research Program through a gift from Upstage Lung Cancer, the Koch Institute’s Marble Center for Cancer Nanomedicine, and Johnson & Johnson. A.P.S. acknowledges support from the NIH Molecular Biophysics Training Grant and the National Science Foundation Graduate Research Fellowship. J.D.K. acknowledges fellowship support from the Ludwig Center at MIT’s Koch Institute. C.S.W. acknowledges support from the National Science Foundation Graduate Research Fellowship. C.M.C. acknowledges support from the NIH Pre-Doctoral Training Grant T32GM007287. S.N.B. is a Howard Hughes Medical Institute Investigator.

## Author contributions

Conceptualization: A.P.S., J.D.K., S.N.B.; Methodology: A.P.S., J.D.K., A.M.J.; Software: A.P.S.; Validation: A.P.S., J.D.K., C.S.W., A.M.J.; Formal analysis: A.P.S., J.D.K., C.S.W.; Investigation: A.P.S., J.D.K., C.S.W., A.M.J., S.S., Q.Z.; Resources: A.P.S., J.D.K., A.M.J., S.N., C.M.C., T.J., S.N.B.; Data curation: A.P.S., J.D.K.; Writing–original draft: A.P.S., J.D.K., S.N.B.; Writing–review and editing: A.P.S., J.D.K., C.S.W., A.M.J., S.S., S.N., Q.Z., C.M.C., T.J., S.N.B.; Visualization: A.P.S.; Supervision: A.P.S., J.D.K., T.J., S.N.B.; Project administration: A.P.S., J.D.K., S.N.B.; Funding acquisition: A.P.S., J.D.K., S.N., T.J., S.N.B.

## Competing interest statement

T.J. is a member of the Board of Directors of Amgen and Thermo Fisher Scientific and a co-founder of Dragonfly Therapeutics and T2 Biosystems; serves on the Scientific Advisory Board of Dragonfly Therapeutics, SQZ Biotech, and Skyhawk Therapeutics; and is President of Break Through Cancer. T.J.’s laboratory currently receives funding from Johnson & Johnson and The Lustgarten Foundation, but these funds did not support the research described in this manuscript. S.N.B. is a director at Vertex; is a co-founder of and consultant for Glympse Bio, Satellite Bio, and Cend Therapeutics; holds equity in Glympse Bio, Satellite Bio, Cend Therapeutics, and Catalio Capital; consults for Moderna; and receives sponsored research funding from Johnson & Johnson. Funds from Johnson & Johnson to S.N.B.’s laboratory supported the research described in this manuscript.

