## Supplementary Information for "Multiscale profiling of enzyme activity in cancer"

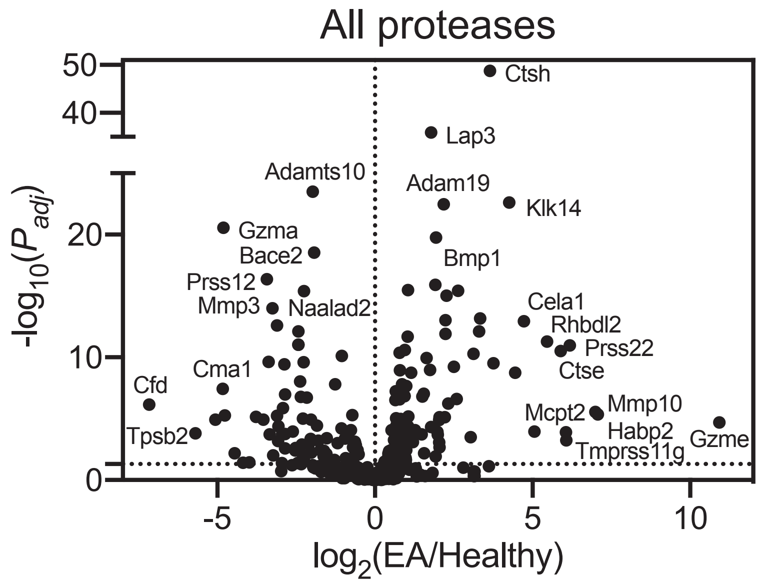


Figure S1: **Proteases are differentially expressed in the Eml4-Alk mouse model of NSCLC.** An existing bulk RNA-seq dataset from the Eml4-Alk model [24] was analyzed to identify endoproteases differentially expressed between Eml4-Alk mice (EA) and healthy controls (Healthy). Each point represents one protease gene. Significance was calculated by Wald test followed by adjustment for multiple hypotheses using the Benjamini-Hochberg correction. Dotted line is at $P_{adj} = 0.05$.

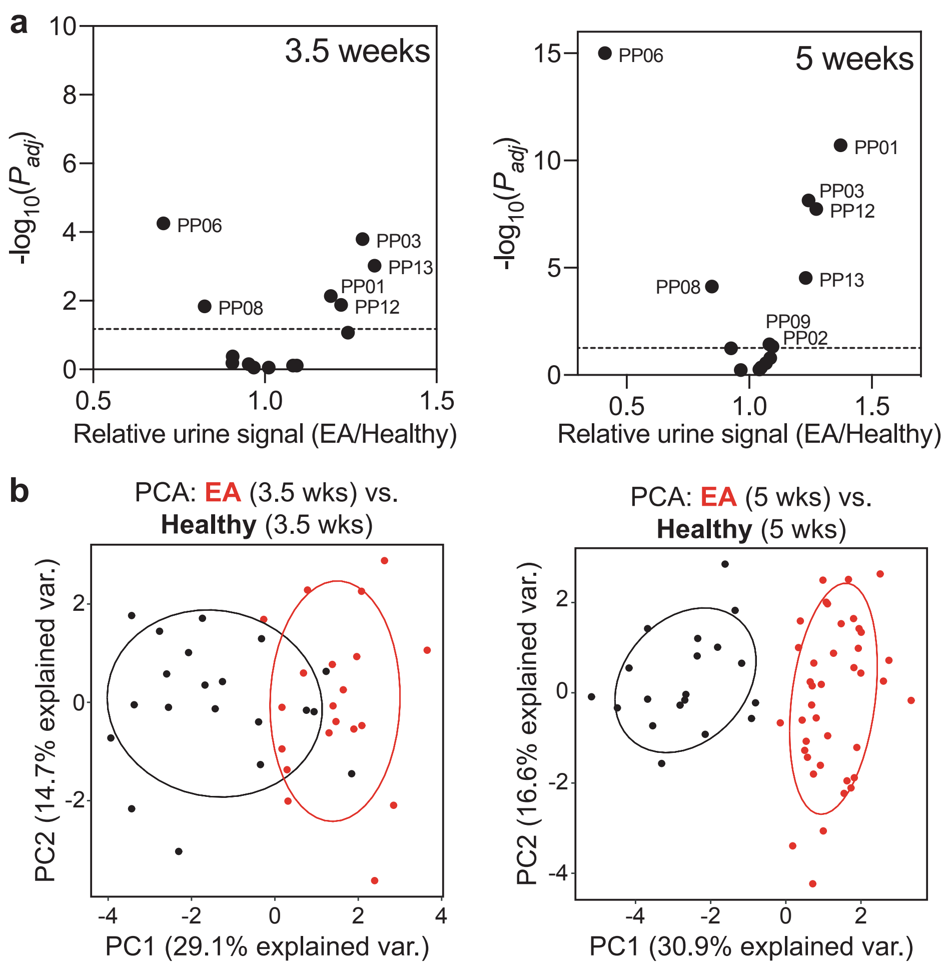


Figure S2: **Activity-based nanosensors differentiate mice bearing ALK$^+$ NSCLC from healthy controls. (a)** Mean scaled urinary reporter concentrations in Eml4-Alk (EA) and healthy (Healthy) mice were compared at 3.5 weeks (n = 20 per group; left) and 5 weeks (EA, n = 40; Control, n = 19; right) after tumor induction, and $-\log_{10}(P_{adj})$ was plotted against fold change between EA and healthy mice. Significance was calculated by two-tailed t-test with Holm-Sidak correction. Dotted line is at $P_{adj} = 0.05$. **(b)** PCA of urinary reporter output of EA mice and healthy controls at 3.5 weeks (n = 20 per group; left) and 5 weeks (EA, n = 40; Healthy, n = 19; right) after tumor induction.

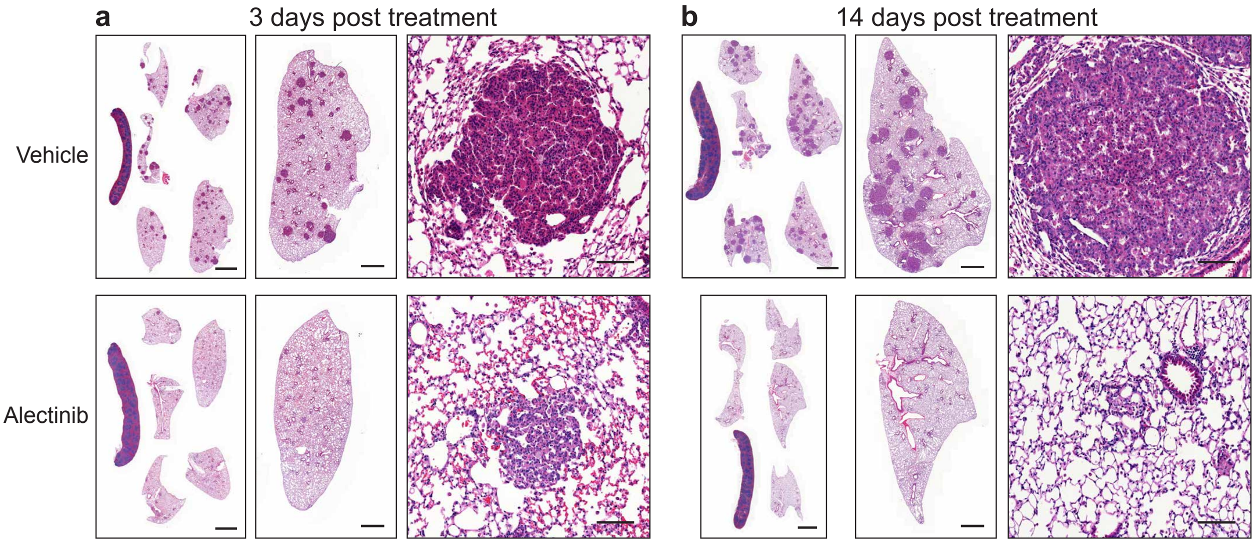


Figure S3: **Alectinib treatment induces tumor regression in the Eml4-Alk model. (a, b)** Images of representative lung sections and tumors from vehicle-treated and alectinib treated-mice, 3 days (a) and 14 days (b) post treatment initiation. Scale bars = 2 mm, 1 mm, 100 $\mu$m for all lobes, single lobe, and individual tumors, respectively.

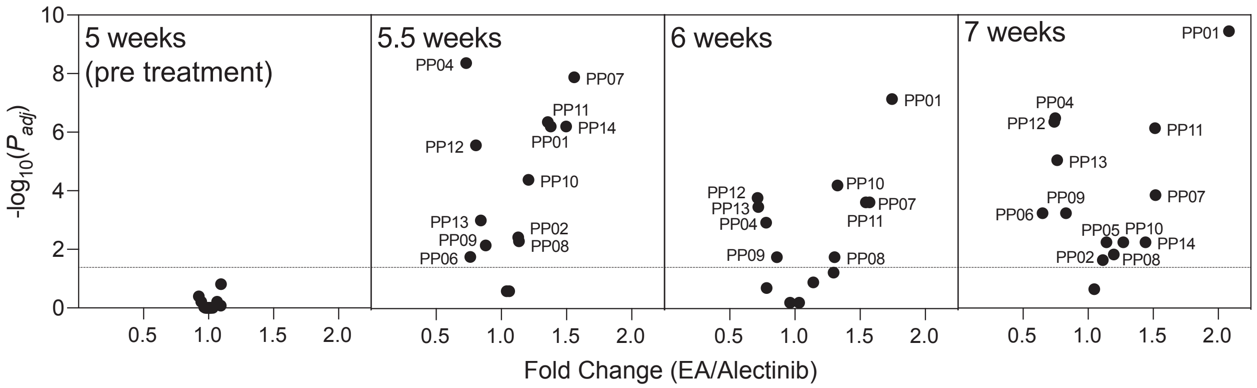


Figure S4: **Multiple reporters are differentially enriched in the urine of vehicle-treated and alectinib-treated Eml4-Alk mice.** Mean scaled urinary reporter concentrations in vehicle-treated (EA) and alectinib-treated (Alectinib) Eml4-Alk mice were compared at 5 weeks (pre-treatment; EA, n = 20; Alectinib, n = 20), 5.5 weeks (EA, n = 19; Alectinib, n = 19), 6 weeks (EA, n = 13; Alectinib, n = 12), and 7 weeks (EA, n = 14; Alectinib, n = 14) after tumor induction, with alectinib/vehicle treatment beginning at 5 weeks post tumor induction. $-\log_{10}(P_{adj})$ was plotted against fold change between vehicle- and alectinib-treated Eml4-Alk mice. Significance was calculated by two-tailed t-test followed by adjustment for multiple hypotheses with Holm-Sidak correction. Dotted line is at $P_{adj} = 0.05$.

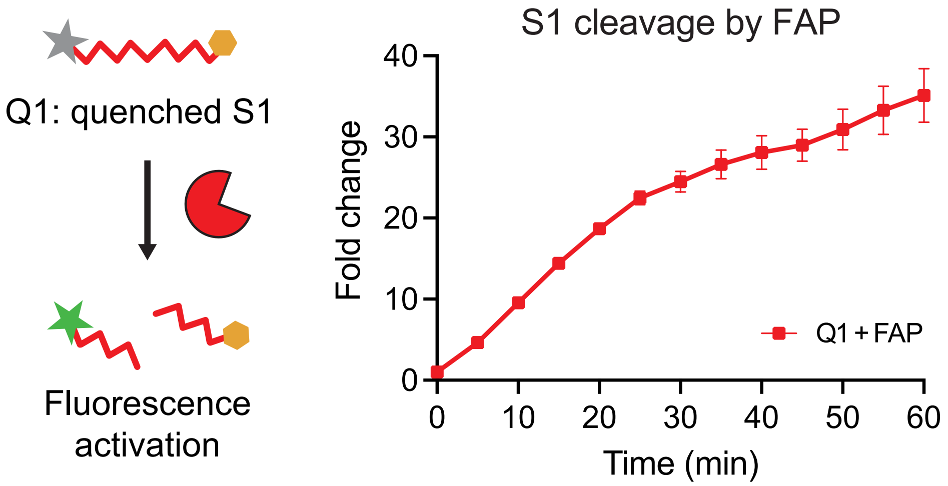


Figure S5: **Cleavage of S1 by recombinant fibroblast activation protein (FAP).** The quenched probe Q1, incorporating the substrate S1, was incubated with recombinant FAP, and fluorescence activation was monitored over time (n = 3 replicates; mean ± s.d.).

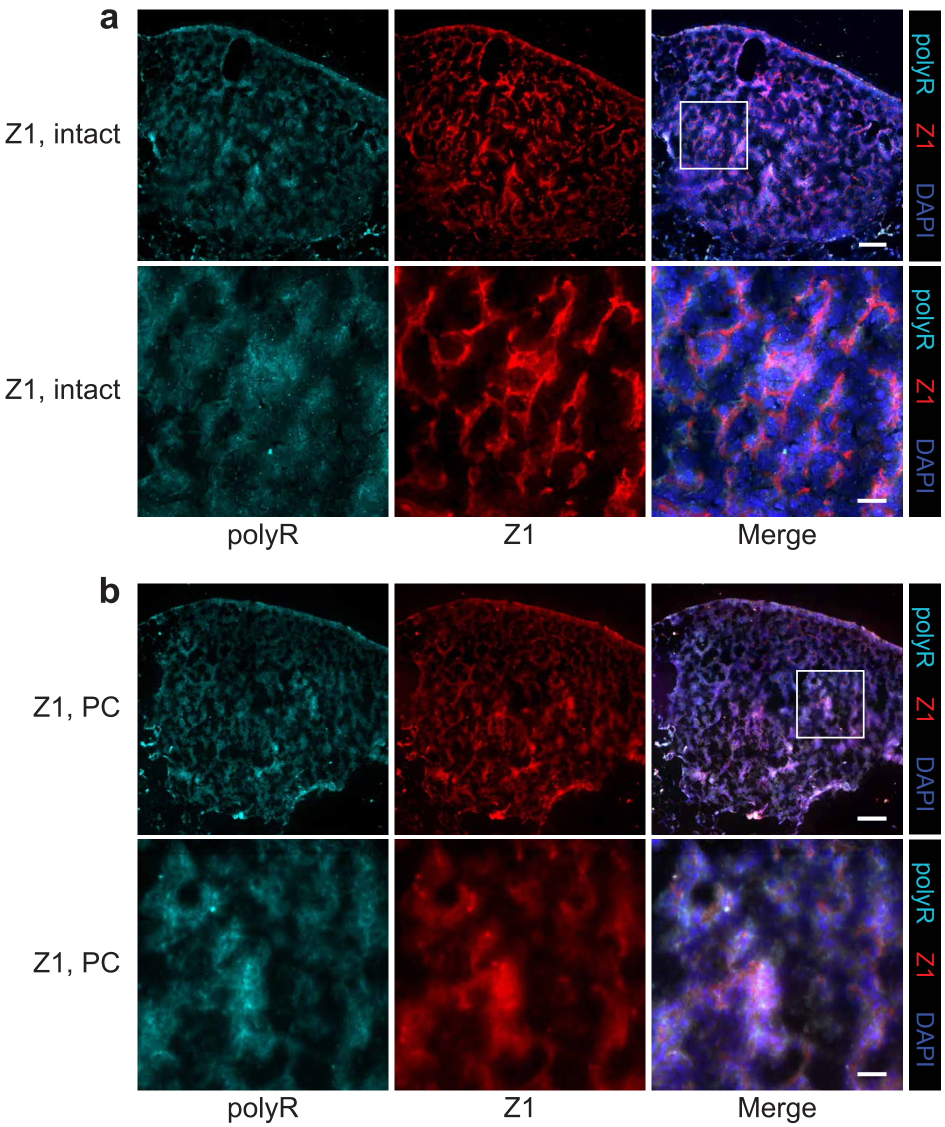


Figure S6: **Z1 localization requires activation by endogenous, tumor-resident proteases. (a, b)** Tissue labeling and localization of intact (a) or FAP-precleaved Z1 (PC; b), together with free polyR control (cyan), within Eml4-Alk tumors. Probes were incubated at 37° C for 4 hours to allow cleavage of intact Z1 by endogenous, tissue-resident proteases. Scale bars = 100 $\mu$m, 25 $\mu$m (lower and higher magnification, respectively).

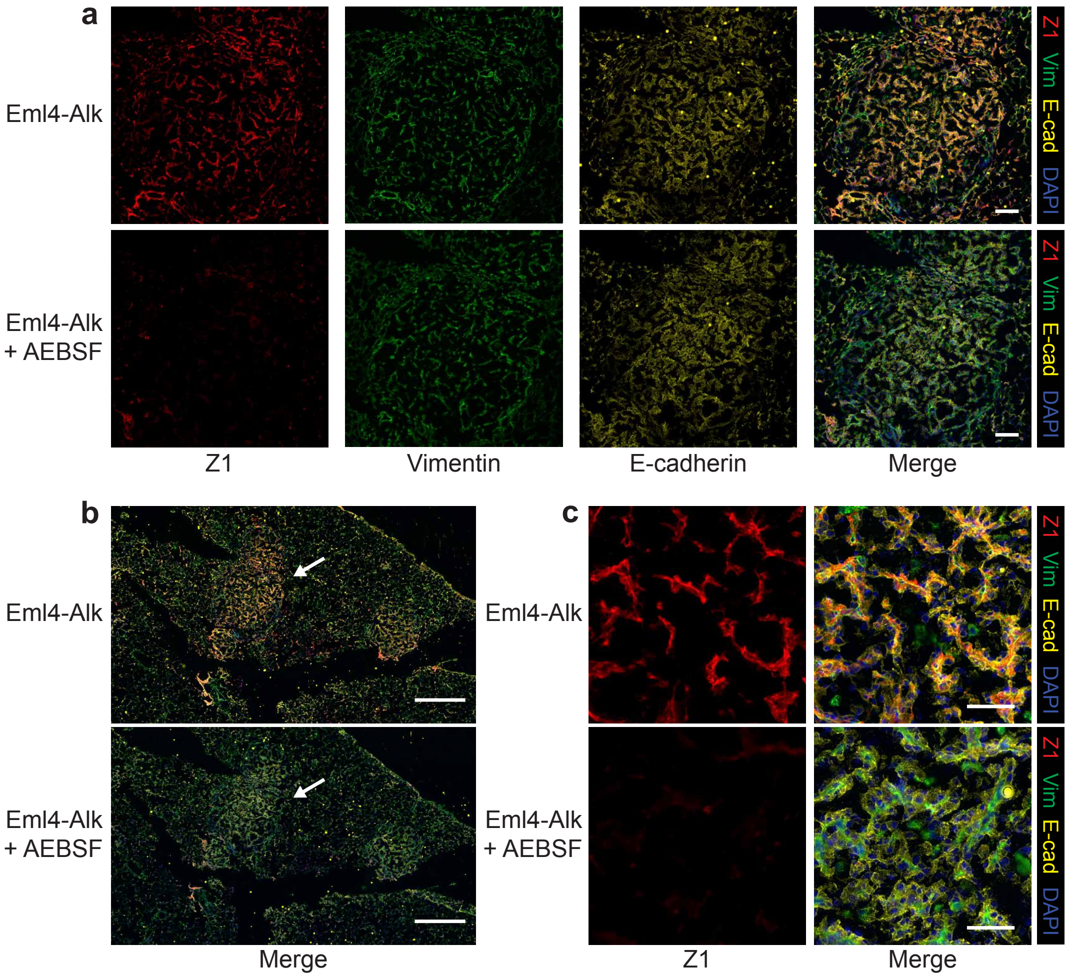


Figure S7: **Z1, vimentin, and E-cadherin staining in Eml4-Alk tissue.** **(a)** Staining of representative Eml4-Alk tumor with Z1 (red), together with the mesenchymal marker vimentin (green) and the epithelial marker E-cadherin (yellow), with or without the serine protease inhibitor AEBSF. Scale bars = 100 $\mu$m. **(b)** Z1, vimentin, and E-cadherin staining across Eml4-Alk lungs. Scale bars = 500 $\mu$m. **(c)** Higher-magnification images of stained tissue, with or without AEBSF, in representative tumor regions indicated by arrows in (b). Scale bars = 50 $\mu$m.

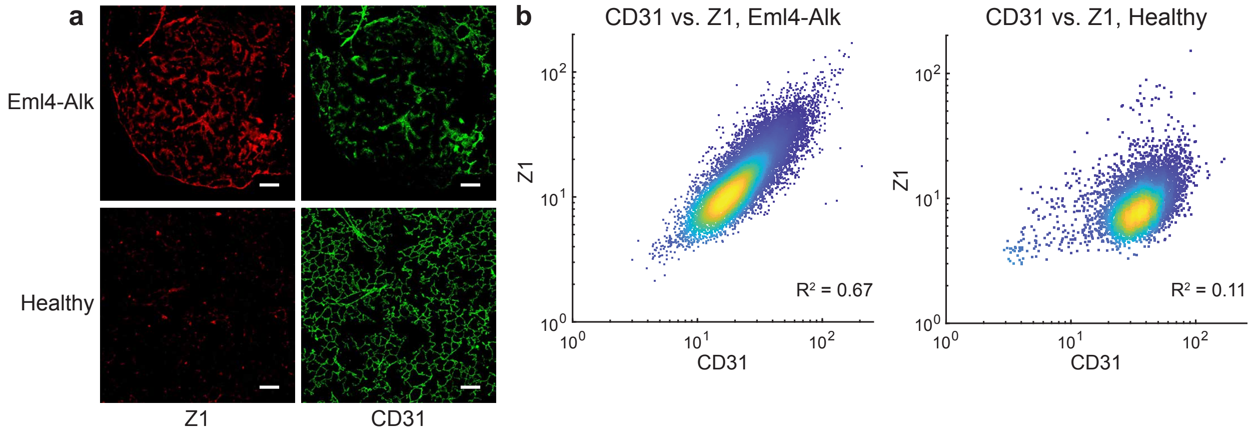


Figure S8: **Z1 and CD31 staining in Eml4-Alk and healthy lungs.** **(a)** Application of Z1 (red) to lung tissue sections from Eml4-Alk and healthy control mice, together with the endothelial marker CD31 (green). Scale bars = 100 $\mu$m. **(b)** Cell-by-cell quantification and correlation of Z1 and CD31 fluorescence intensity in Eml4-Alk (left) and healthy (right) lung tissue sections ($R^2 = 0.67, R^2 = 0.11$ for Eml4-Alk and healthy, respectively).

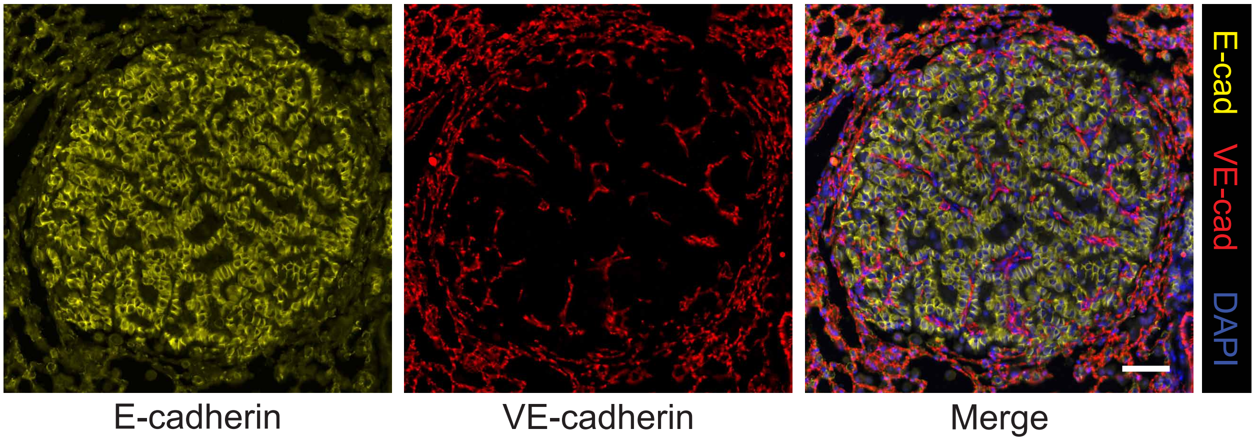


Figure S9: **VE-cadherin expression in Eml4-Alk tumors.** Immunofluorescence staining for E-cadherin (yellow) and the endothelial marker VE-cadherin (red) in formalin-fixed, paraffin-embedded Eml4-Alk lung tissue sections, with images from a representative tumor shown. Sections were counterstained with DAPI (blue). Scale bar = 50 $\mu$m.

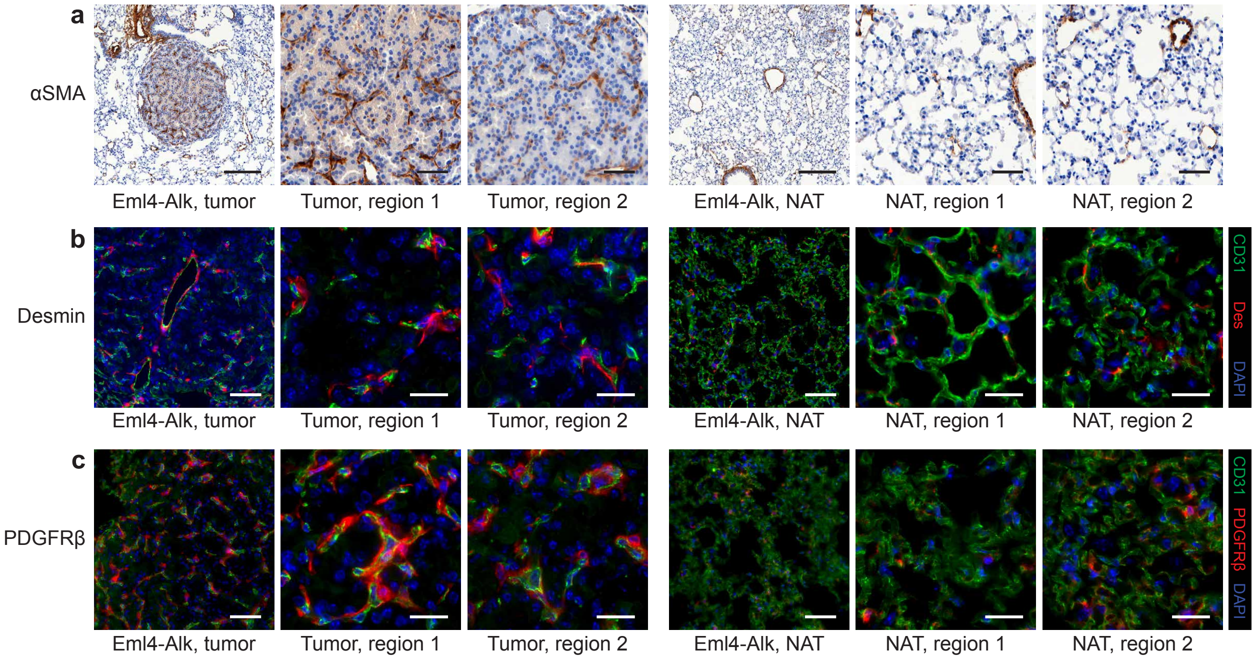


Figure S10: **Abundance of pericyte markers in Eml4-Alk tumors and normal adjacent tissue. (a)** Immunohistochemical staining for the smooth muscle and pericyte marker $\alpha$-smooth muscle actin ($\alpha$SMA, C) in representative Eml4-Alk tumor and normal adjacent tissue (NAT) regions. Scale bars = 200 $\mu$m (left columns per group), 50 $\mu$m (center and right columns per group). **(b, c)** Immunofluorescence staining for the endothelial cell marker CD31 (green) with each of the pericyte markers desmin (red, b) and PDGFR$\beta$ (red, c) in representative Eml4-Alk tumor and NAT regions. Sections were counterstained with DAPI (blue). Scale bars = 50 $\mu$m (left columns per group), 20 $\mu$m (center and right columns per group).

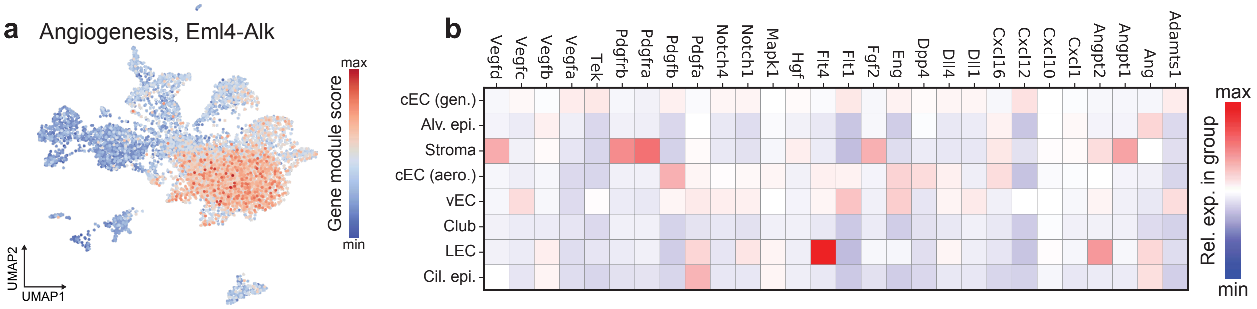


Figure S11: **Expression of angiogenesis module across the cellular landscape of Eml4-Alk lungs.** **(a)** Collective expression score for angiogenesis gene module mapped onto UMAP of cells from Eml4-Alk lungs. **(b)** Relative expression levels of individual genes in angiogenesis module, standardized across cell type clusters in Eml4-Alk lungs. Matrix plot is colored by mean expression values for a given gene, averaged across all cells in a cluster.

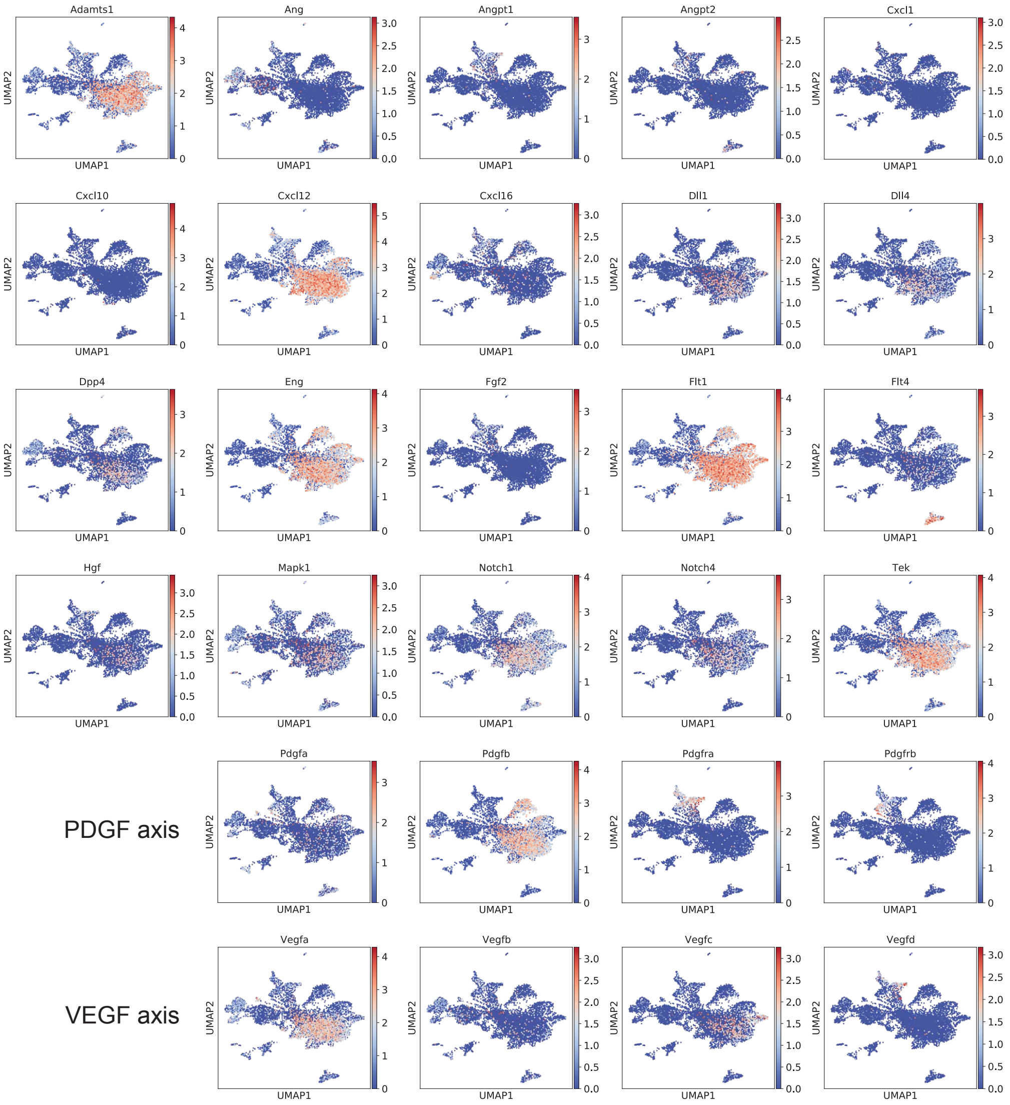


Figure S12: **Expression levels of angiogenesis module genes across the cellular landscape of Eml4-Alk lungs.** Expression of individual genes in the angiogenesis module (Fig. S11) was mapped onto the UMAP of cells from Eml4-Alk lungs.

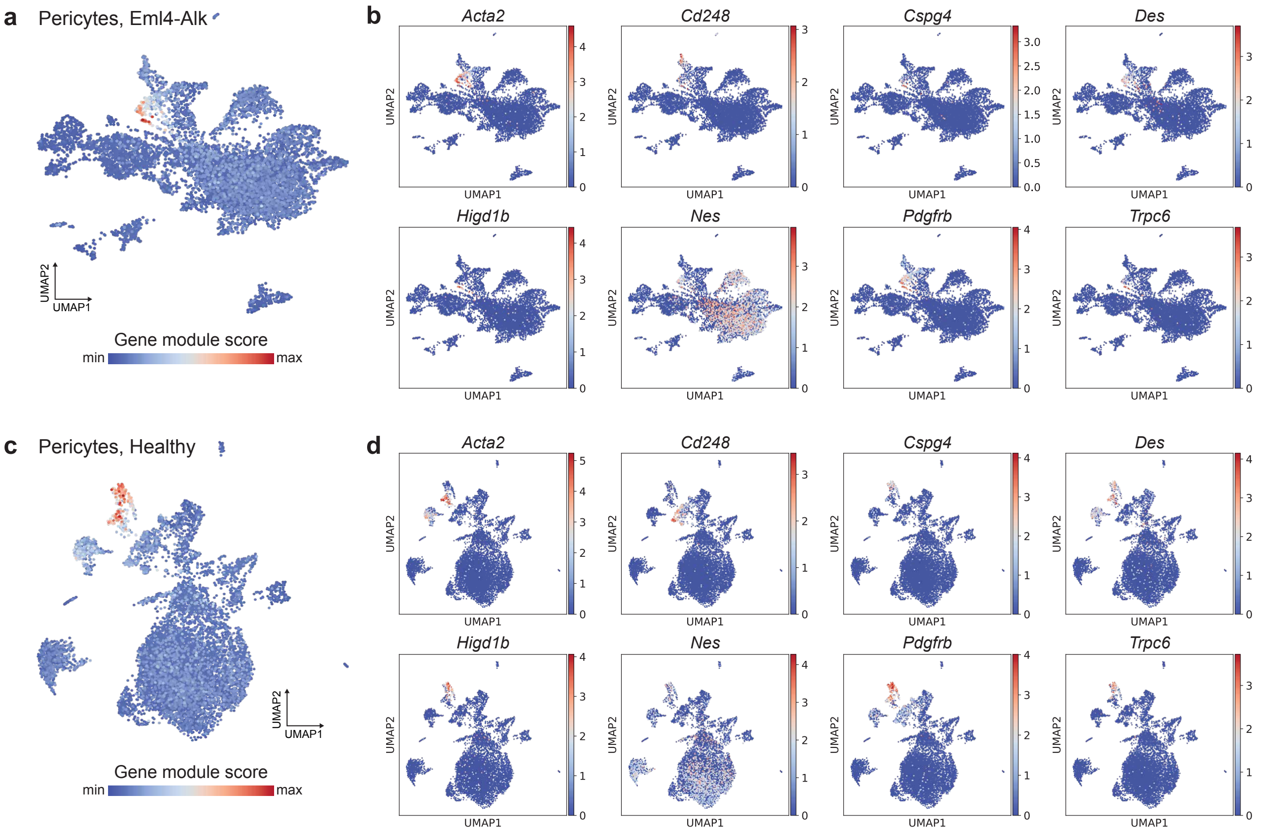


Figure S13: **Distribution of pericyte marker genes in Eml4-Alk and healthy lungs.** **(a)** Collective expression score for pericyte marker gene module mapped onto the UMAP of cells from Eml4-Alk lungs. **(b)** Relative expression levels of individual pericyte marker genes mapped onto the UMAP of cells from Eml4-Alk lungs. **(c)** Collective expression score for pericyte marker gene module mapped onto the UMAP of cells from healthy lungs. **(d)** Relative expression levels of individual pericyte marker genes mapped onto the UMAP of cells from healthy lungs.

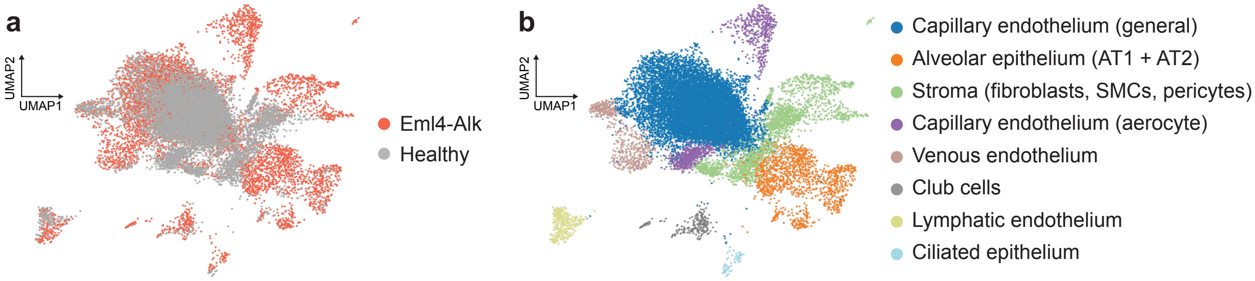


Figure S14: **Integrated comparison of single cell transcriptomic landscapes of Eml4-Alk and healthy lungs. (a, b)** UMAP of integrated dataset of single cells from Eml4-Alk and healthy lungs (pooled samples of n = 3 mice per condition). UMAP is colored by condition of origin (a) or inferred cell type (b). Individual dots correspond to individual cells.

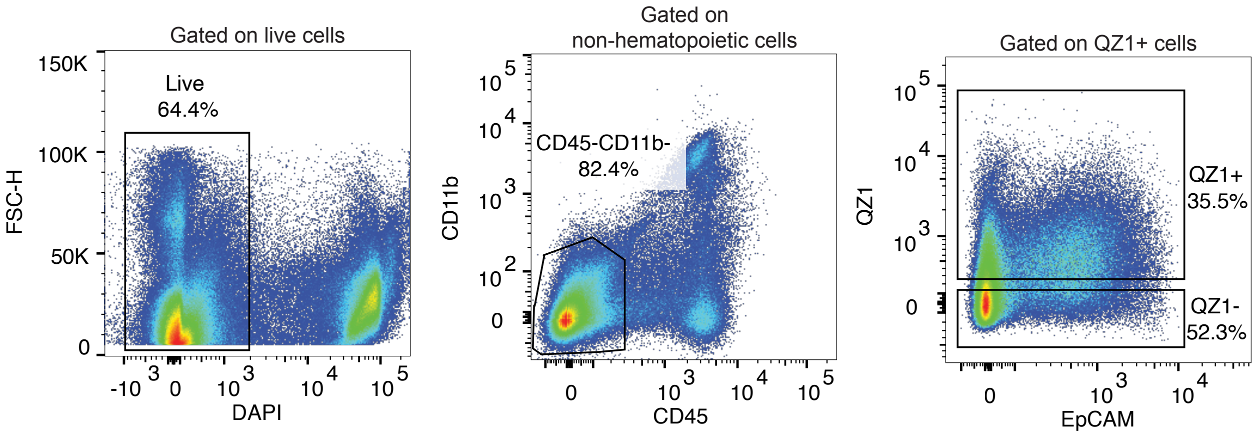


Figure S15: **Gating strategy for activity-based cell sorting.** For activity-based cell sorting on the basis of QZ1 signal, cells were first sorted on the basis of viability by gating on DAPI signal. All non-hematopoietic cells were isolated by gating on CD45 and CD11b negativity. Finally, cells were sorted for signal from the QZ1 activity probe by gating signal against EpCAM.

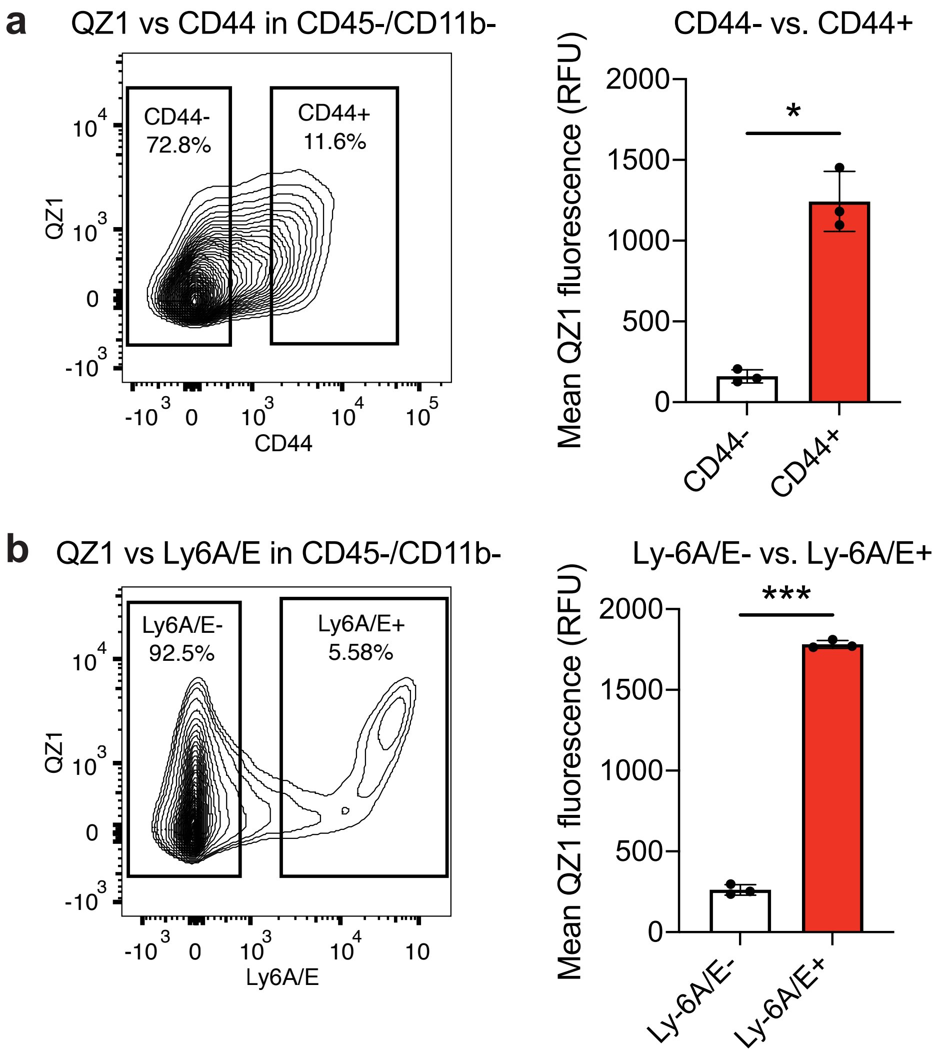


Figure S16: **Concurrent flow cytometry analysis of activity probe signal and cell surface markers. (a, b)** Flow cytometry plot (left) and quantification (right) of Cy5 (QZ1) fluorescence intensity in CD45-, CD11b- cells from Eml4-Alk lungs, gated by the cell surface markers CD44 (a) and Ly-6A/E (b) (n = 3 biological replicates; mean ± s.d.; two-tailed paired t-test, $*P = 0.0120$ for CD44, $***P = 0.000374$ for Ly-6A/E).

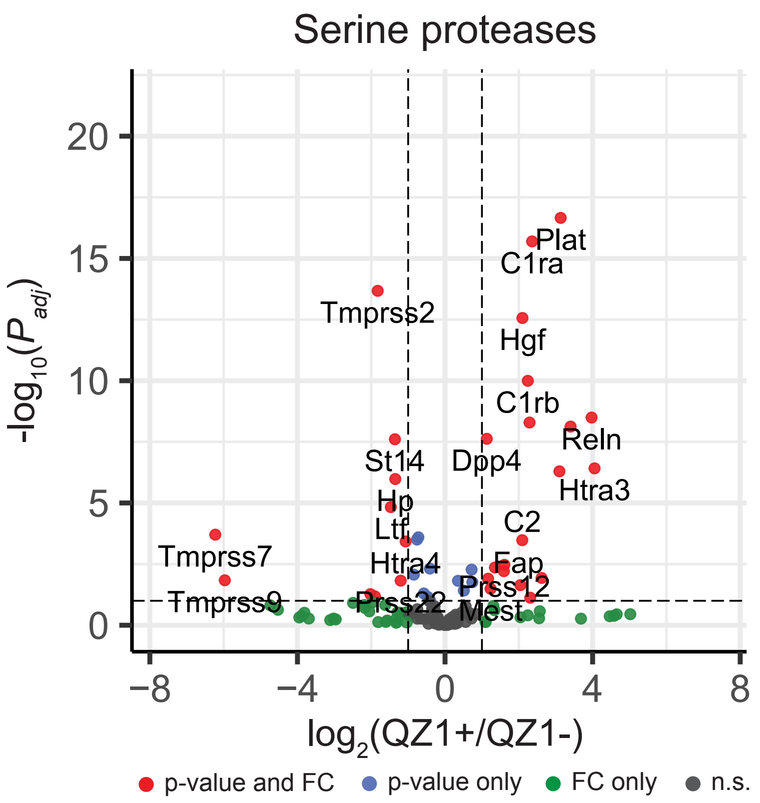


Figure S17: **Serine proteases are differentially expressed with respect to QZ1 label.** Differential gene expression analysis, filtered for serine proteases, comparing the QZ1+ and QZ1- sorted populations from Eml4-Alk lungs. Each point represents one protease gene. Significance was calculated by Wald test followed by adjustment for multiple hypotheses using the Benjamini-Hochberg correction. Dotted line is at $P_{adj} = 0.01$.

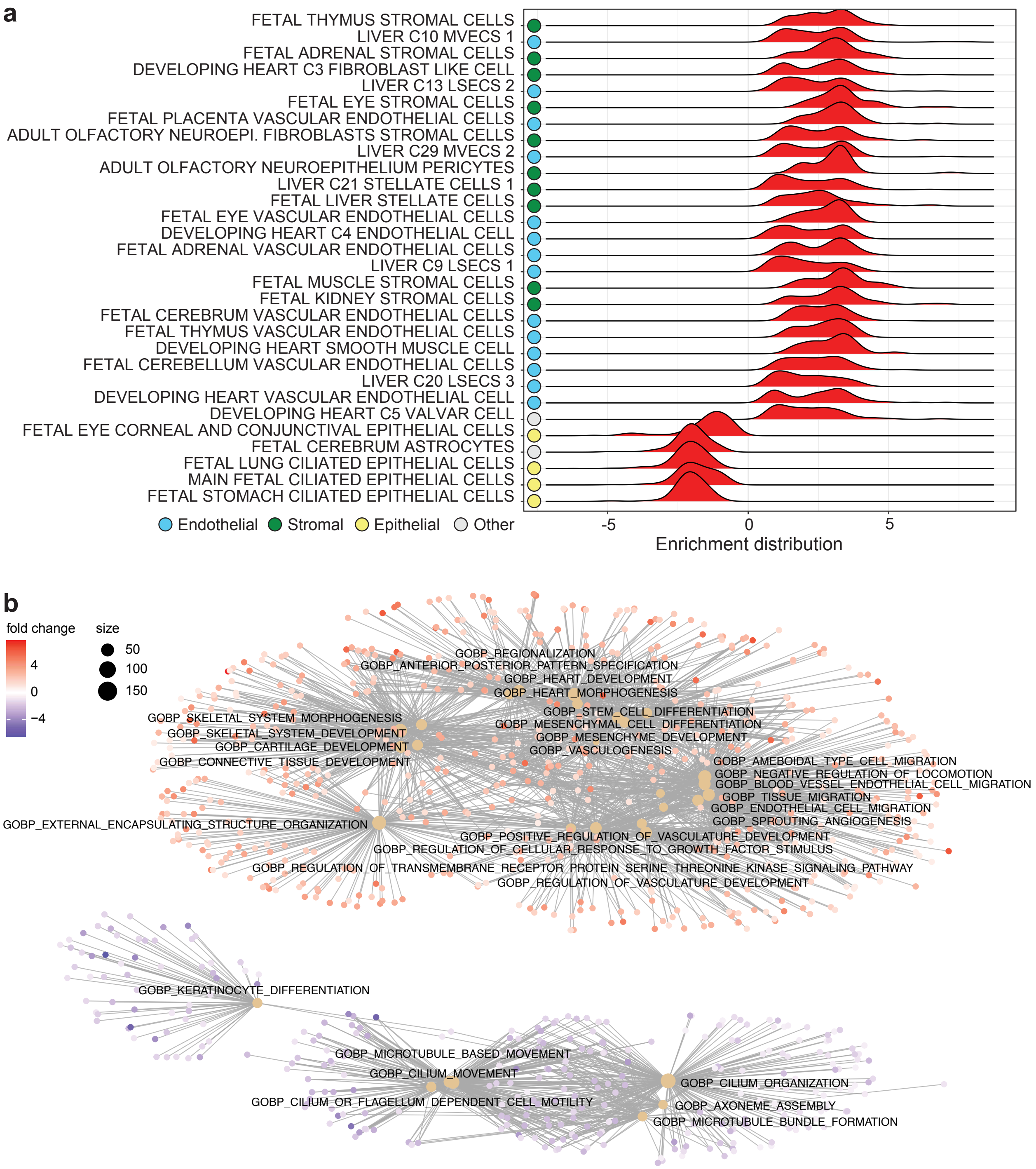


Figure S18: **Gene set enrichment analysis downstream of activity-based cell sort.** **(a)** Gene set enrichment analysis for cell type-specific expression modules based on a ranked list of differentially expressed genes from QZ1+ and QZ1- populations isolated via activity-based cell sorting. **(b)** Enrichment map of functional gene ontology (GO) modules from Fig. 7G. Mutually overlapping gene sets are clustered together, while non-overlapping gene sets are distanced and clustered separately.

| Name | Reporter | Photolabile group | Substrate | Nanocarrier |
|---|---|---|---|---|
| PP01 | e(+2G)(+6V)ndneeGFFsAr | ANP | GGPQGIWGQC | PEG-$8_{40kDa}$ |
| PP02 | eG(+6V)ndneeGF(+1F)s(+1A)r | ANP | GGPVGLIGC | PEG-$8_{40kDa}$ |
| PP03 | e(+3G)(+1V)ndneeGFFs(+4A)r | ANP | GGPVPLSLVMC | PEG-$8_{40kDa}$ |
| PP04 | e(+2G)Vndnee(+2G)FFs(+4A)r | ANP | GGPLGLRSWC | PEG-$8_{40kDa}$ |
| PP05 | eGVndnee(+3G)(+1F)Fs(+4A)r | ANP | GGPLGVRGKC | PEG-$8_{40kDa}$ |
| PP06 | e(+2G)(+6V)ndnee(+3G)(+1F)(+1F)s(+1A)r | ANP | GGfPRSGGGC | PEG-$8_{40kDa}$ |
| PP07 | eG(+6V)ndnee(+3G)(+1F)Fs(+4A)r | ANP | GGLGPKGQTGC | PEG-$8_{40kDa}$ |
| PP08 | e(+3G)(+1V)ndneeG(+10F)FsAr | ANP | GGGSGRSANAKGC | PEG-$8_{40kDa}$ |
| PP09 | eGVndneeGF(+10F)s(+4A)r | ANP | GGKPISLISSGC | PEG-$8_{40kDa}$ |
| PP10 | e(+2G)(+6V)ndneeG(+10F)(+1F)s(+1A)r | ANP | GGILSRIVGGGC | PEG-$8_{40kDa}$ |
| PP11 | e(+3G)(+1V)ndnee(+2G)(+10F)Fs(+4A)r | ANP | GGSGSKIIGGGC | PEG-$8_{40kDa}$ |
| PP12 | eGVndneeG(+10F)(+10F)sAr | ANP | GGPLGMRGGC | PEG-$8_{40kDa}$ |
| PP13 | e(+2G)(+6V)ndnee(+3G)(+10F)(+1F)s(+4A)r | ANP | GGP-(Cha)-G-Cys(Me)-HAGC | PEG-$8_{40kDa}$ |
| PP14 | e(+3G)(+1V)ndnee(+2G)(+10F)(+10F)sAr | ANP | GGAPFEMSAGC | PEG-$8_{40kDa}$ |

Table S1: **Reporter and substrate sequences for *in vivo* activity-based nanosensors.** Lowercase letters: d-amino acids; ANP: 3-Amino-3-(2-nitro-phenyl)propionic acid; Cha: 3-Cyclohexylalanine; Cys(Me): methyl-cysteine

| Name | Sequence | Readout |
|---|---|---|
| Z1 | UeeeeeeeeXGGPQGIWGQGrrrrrrrrrX-k(Cy5) | Fluorescence (in situ) |
| Z7 | UeeeeeeeeXGGLGPKGQTGGrrrrrrrrrX-k(Cy3) | Fluorescence (in situ) |
| Z10 | UeeeeeeeeXGILSRIVGGGrrrrrrrrrX-k(FAM) | Fluorescence (in situ) |
| QZ1 | (QSY21)-eeeeeeeee-c(PEG2K)-oGGPQGIWGQG-rrrrrrrrr-k(Cy5) | Fluorescence (in vitro / in vivo) |
| polyR | rrrrrrrrrX-k(Cy7) | Fluorescence (in situ) |
| S1 | GGPQGIWGQC | Cleavage motif |

Table S2: **Peptide sequences for activatable zymography probes.** Lowercase letters: d-amino acids; QSY21-Cy5: FRET pair, with Cy5 as fluorophore and QSY21 as quencher; PEG2K: (poly)ethylene-glycol, MW 2000 g/mol; o: 5-amino-3-oxopentanoyl; U: succinoyl; X: 6-aminohexanoyl

| | Cap. endothelium (general) | | Stroma | | Cap. endothelium (aerocyte) | | Ciliated epithelium | |
|---|---|---|---|---|---|---|---|---|
| **Gene** | **$\log_2$(EA/WT)** | **$P_{adj}$** | **$\log_2$(EA/WT)** | **$P_{adj}$** | **$\log_2$(EA/WT)** | **$P_{adj}$** | **$\log_2$(EA/WT)** | **$P_{adj}$** |
| *Pdgfa* | 0.578 | $8.29 \times 10^{-1}$ | 0.07840 | 1.00 | 1.670 | $3.85 \times 10^{-1}$ | 0.999 | 1.00 |
| *Pdgfb* | 0.676 | $1.39 \times 10^{-25}$ | 0.824 | 1.00 | 0.835 | $1.77 \times 10^{-11}$ | 10.331 | 1.00 |
| *Pdgfra* | -0.396 | 1.00 | -0.0295 | 1.00 | -0.145 | 1.00 | 11.335 | 1.00 |
| *Pdgfrb* | -1.776 | 1.00 | 0.178 | 1.00 | -7.183 | 1.00 | 0.000 | 1.00 |
| *Cxcl12* | 1.453 | $1.73 \times 10^{-195}$ | 1.542 | $5.77 \times 10^{-19}$ | 1.537 | $1.32 \times 10^{-2}$ | 11.335 | 1.00 |

Table S3: **Differential expression analysis of PDGF signaling genes in scRNA-seq data from Eml4-Alk and healthy lungs.** Differential gene expression analysis was conducted on scRNA-seq data from cells within each of the indicated cell compartments, and results were filtered for genes from the PDGF signaling axis. Significance was calculated by the Wilcoxon rank-sum test with Benjamini-Hochberg correction.
